# Bone microarchitecture and material properties decline differently across midlife for male and female F344 × BN F1 rats

**DOI:** 10.64898/2026.04.16.719016

**Authors:** Hunter Hasskamp, Emelia Keim, Kenna Brown, Sarah Sucher, Chelsea M. Heveran, Stephen A. Martin

**Author notes:** Corresponding Author, 2155 Analysis Dr. Bozeman, MT, USA 59718. Senior Authors contributed equally (Chelsea M. Heveran, Stephen A. Martin).

## Abstract

While bone mineral density (BMD) remains the clinical standard for assessing age-related fracture risk, accumulating evidence indicates that bone quality, including matrix properties and microarchitecture, contributes to fracture susceptibility in ways not captured by BMD alone. As matrix-targeted therapeutics emerge, preclinical models that exhibit translationally relevant bone quality changes are needed. Here, we evaluated the Fischer 344 × Brown Norway (F344×BN) F1 rat, a strain characterized by hybrid vigor and non-pathological aging, as a model for studying matrix-related mechanisms of skeletal aging. Femurs from male and female rats aged 7, 15, and 22 months were analyzed to quantify age- and sex-dependent changes in bone microarchitecture, fracture resistance, and matrix properties. Microcomputed tomography analyses revealed sexually dimorphic aging trajectories. From 7 to 22 months, females exhibited moderate declines in trabecular microarchitecture and no change in cortical porosity, whereas males showed pronounced trabecular deterioration and increased cortical porosity. Whole-bone flexural testing demonstrated age-related declines in material properties that were not attributable to changes in geometry, while females maintained geometry-scaled bone strength. Both sexes exhibited reduced bone toughness with age. Raman spectroscopy identified matrix-level alterations in males by 15 months, whereas systemic markers of bone turnover remained unchanged across age or sex. Together, these findings indicate that males exhibit combined tissue-scale and whole-bone deterioration by midlife, while females exhibit declining fracture resistance preceding substantial cortical bone loss or overt matrix deterioration. These results support the F344×BN F1 rat as a translational model for investigating matrix-driven skeletal aging.

**Lay summary:** F344 x BN F1 hybrid rats provide a healthy, matrix-driven skeletal aging model. This strain exhibits distinct aging trajectories dependent on sex. Strength and toughness decrease in both sexes by midlife. Fracture resistance declines in females prior to substantial bone loss.

## 1.0 Introduction

Skeletal aging is a multifactorial process involving progressive changes in bone mass, structure, and material properties that collectively increase fracture risk. Aging-related fractures are highly prevalent and constitute a major public health burden, contributing to loss of independence, diminished quality of life, and increased mortality. Globally, fractures account for billions of dollars in annual healthcare costs, largely driven by surgical repair, rehabilitation, and long-term care [1]. In response, numerous therapeutic strategies and clinical indices have been developed to mitigate these issues. However, bone mineral density remains the primary clinical measure of skeletal health, despite its limitations in capturing age-related declines in bone quality that influence fracture susceptibility [2]. Of note, clinical definitions of osteoporosis only capture ∼50% of hip fractures in women aged 65+ [3]. To address this issue, interventions that target bone matrix are beginning to emerge in preclinical studies [4]. The development of such interventions first requires well-characterized models that exhibit clinically relevant, age-related changes in bone quality.

For maximal translational relevance, skeletal aging should be evaluated during midlife before the onset of fragility fractures. For postmenopausal women in the United States, the medical costs of bone fragility (direct medical costs and indirect costs related to productivity loss and caregiving) were $57 billion in 2018 and are expected to rise to $95 billion by 2040 [1]. Intervening at this stage has the potential to reduce both the financial burden and decline in quality of life associated with fracture repair and rehabilitation in older age. Rodent models are central to skeletal aging research, providing insight into bone development, metabolism, genetic regulation, and repair. Mice are commonly used due to their low cost, shorter lifespan, their ability to be genetically modified, and have been extensively characterized for age-related bone loss and fracture resistance decline [5]. Across rodent species, aging is associated with trabecular bone loss, preservation of apparent whole-bone strength through architectural remodeling, changes in mineral and collagen properties, and reduced fracture resistance. However, rodent models do not fully recapitulate all features of human skeletal aging, and strain- and species-specific differences can limit reproducibility and translational relevance. For example, C57BL/6 mice exhibit bone loss accompanied by reduced matrix quality, strength, and toughness by midlife, paralleling aspects of human skeletal aging, but these changes are not solely age-driven [6,7]. When housed at standard room temperature (∼22 °C), C57BL/6 mice experience premature bone loss driven by elevated sympathetic activity associated with non-shivering thermogenesis [8]. Although housing at thermoneutrality (∼30 °C) prevents this phenotype, long-term maintenance under these conditions is logistically challenging and costly [8]. In contrast, rats do not experience housing-induced bone loss and more closely approximate human skeletal physiology due to their larger size, homeothermic regulation, and slower rate of age-related skeletal decline [9]. Accordingly, both outbred (e.g., Wistar, Sprague-Dawley (SD)) and inbred (Fischer 344 (F344)) rat strains have been used in skeletal aging research, though findings across strains remain variable and partially concordant with human skeletal aging in terms of trabecular bone loss, development of cortical porosity, and declining material properties with age [10–15]. Together, these limitations highlight the need for aging models that isolate intrinsic skeletal aging during midlife and better reflect the physiology of the aging human skeleton.

The Fischer 344 × Brown Norwegian (F344 × BN) F1 hybrid rat is widely used in aging research outside of the bone research community due to its hybrid vigor, which confers extended lifespan, reduced incidence of spontaneous disease, delayed onset of age-related pathologies, and gradual phenotypic aging across multiple organ systems [16]. This strain has been extensively utilized to study metabolism, cardiovascular function, cognition, and responses to caloric restriction, and serves as a reference model in aging research [e.g.,17-20]. Despite this widespread use, age- and sex-dependent skeletal trajectories in F344 × BN F1 rats remain poorly characterized. In particular, it remains unclear how fracture resistance evolves across adulthood and midlife in this strain. In humans, middle age is a critical period where bone quality declines and preventive interventions are likely to have the greatest impact. It is also essential to include both sexes in this research as sex is an essential biological variable for aging and preclinical research [21]. This gap limits the ability to interpret prior findings, compare interventions across studies, and design therapies that target bone quality rather than bone mass alone.

To determine matrix changes during skeletal aging in F344 × BN F1 rats, the present study comprehensively characterizes skeletal aging in male and female rats from 7 to 22 months of age, a period analogous to the human midlife [22]. Cortical and trabecular architecture were quantified by microcomputed tomography (µCT); whole-bone mechanical performance was assessed using flexural testing; measured matrix properties were evaluated by Raman spectroscopy; and systemic and local indices of bone turnover were measured. Our findings support the F344 × BN F1 rat as a translationally relevant model for sex-specific skeletal aging and identify midlife as a period in which matrix deterioration and declines in whole-bone fracture resistance emerge, with reductions in whole-bone material properties preceding significant bone loss in females. This model provides a foundation for evaluating interventions aimed at preserving bone matrix quality and reducing fracture risk across the lifespan.

## 2.0 Materials and methods

### 2.1 Animals

The Montana State University Institutional Animal Care and Use Committee approved all animal procedures. Male and female F344 × BN F1 hybrid rats were obtained from the National Institute on Aging Aged Rodent Colony (Charles River Laboratories, Wilmington, MA, USA). A total of 41 animals were used for skeletal analyses, distributed across age and sex: 7-month-old (7-mo) females (n = 8), 7-mo males (n = 8), 15-month-old (15-mo) females (n = 6), 15-mo males (n = 6), 22-month-old (22-mo) females (n = 6), and 22-mo males (n = 7). An additional age- and sex-matched cohort of F344 × BN rats was acquired for serum biomarker analysis supplement: 7-mo females (n = 4), 15-mo females (n = 2), 15-mo males (n = 2), 22-mo females (n = 6), and 22-mo males (n = 1). Rats were housed in groups of three or fewer under controlled conditions (12-hour light/dark cycle) with ad libitum access to water and standard rodent chow (5L79; LabDiet). Animals were acclimated for at least two weeks prior to study procedures.

To label actively mineralizing surfaces, animals were administered two intraperitoneal fluorochrome injections: calcein (20 mg/kg) 10 days prior and alizarin (30 mg/kg) 3 days prior to euthanasia [23]. No adverse events occurred. At the study endpoint, animals were anesthetized with isoflurane and euthanized via exsanguination and cardiac puncture. Femurs and serum were collected. Bones were stripped of soft tissue, wrapped in PBS-soaked gauze, and stored at −20°C. Serum was isolated and stored at −80°C.

### 2.2 Microcomputed tomography

Bone microarchitecture of frozen left femurs in air was assessed using high-resolution microcomputed tomography (μCT40, Scanco Medical AG, Switzerland) using previously established methodologies and guidelines [24]. Scans were performed with an isotropic voxel size of 10 μm^3^, x-ray tube voltage of 70 kVp, current of 114 μA, and 200 ms integration time. Gaussian filtration and global thresholding were applied to segment bone from soft tissue. Trabecular bone was analyzed from a 1.5 mm region beginning 200 μm proximal to the distal growth plate; the region of interest was manually contoured to the endocortical border using a threshold of 310 mgHA/cm³. Measured outcomes included trabecular bone volume fraction (Tb. BV/TV, %), bone mineral density (Tb. BMD, mgHA/cm³), trabecular thickness (Tb. Th, mm), number (Tb. N, mm[¹), separation (Tb. Sp, mm), connectivity density (Conn. D, mm[³), and structure model index (SMI).

Cortical geometry was assessed from 50 transverse slices (10 μm thick) spanning a 500 μm mid-diaphyseal region. A threshold of 700 mgHA/cm³ was applied, and cortical bone was delineated manually. Quantified parameters included total cross-sectional area (Tt. Ar, mm²), cortical area (Ct. Ar, mm²), medullary area (Ma. Ar, mm²), cortical thickness (Ct. Th, mm), bone area fraction (BV/TV, %), cortical porosity (Ct. Po, %), cortical tissue mineral density (Ct. TMD, mgHA/cm³), and cross-sectional moments of inertia (I_max_, I_min_, and pMOI, mm[).

### 2.3 Flexural testing

Whole-bone mechanical properties were assessed using three-point bending with an Instron 5543 load frame (1 kN load cell, USA) and BlueHill 2 software. Left femurs were thawed and hydrated in PBS before testing and placed in a standardized orientation (anterior side down, lateral side outward) on a 16 mm span fixture with rounded contact points. A preload of 5 N was applied, followed by displacement-controlled loading at 3 mm/min until fracture. Load and displacement data were analyzed using a custom MATLAB code (R2024a, MathWorks, USA), integrating bone geometry from μCT to calculate stress-strain behavior using standard equations for bone flexural testing [25]. Outcome measures included stiffness, yield load, ultimate load, energy-to-fracture, elastic modulus, yield stress, and post-yield deformation.

Notched fracture toughness was evaluated on right femora consistent with Bermudez et al. [26]. The proximal and distal ends of femurs were removed. Femurs were notched in the middle of the posterior mid-shaft to a target depth of 1/3 of the anterior-posterior (AP) width using a custom apparatus and saw blade (0.38 mm thickness). Notches were refined by a razor blade coated with 0.05 µm alumina polishing solution. Marrow and alumina solution were flushed using tap water. Notched femora were loaded in three-point bending with the notch facing down until failure with a loading rate of 0.001 mm/sec using the same instrument as above. The span of the three-point bending setup was approximately four times the AP width. The following span definitions were used:

11 mm span if AP width < 3.00 mm

12 mm span if 3.00 ≤ AP width < 3.25 mm

13 mm span if 3.25 ≤ AP width < 3.50 mm

14 mm span if 3.50 ≤ AP width < 3.75 mm

15 mm span if 3.75 ≤ AP width < 4.00 mm

16 mm span if AP width ≥ 4.00 mm

After fracture, the two halves of the diaphysis were centrifuged at 10,000 x g for 5 minutes to remove remaining marrow. The proximal half was submerged in tap water and snap frozen with dry ice for metabolomics. The distal half was frozen in PBS-soaked gauze at −20 °C before being sectioned further. The fracture surface was sectioned off from the distal half with the same saw blade used for notching to be air dried and imaged. The fracture surface was imaged using variable pressure scanning electron microscopy (20 Pa, 15 kV; Zeiss SUPRA 55VP, Germany). Analysis of the fracture surfaces used a custom Python code that measured the cortical bone geometry and the initial notch half-angle. Fracture toughness values were calculated for the K_c,max_ and the K_c,init_ measures. These measures are described in Ritchie et al. [27].

### 2.4 Dynamic histomorphometry

Following μCT and three-point bending analyses, left distal femur segments were embedded in non-infiltrating epoxy (EpoxiCure 2, Buehler, USA). Transverse sections near the mid-diaphysis were cut with a low-speed diamond saw (Smart Cut, Ted Pella, USA) and polished sequentially using 600- and 1000-grit silicon carbide papers, followed by alumina paste in decreasing grit sizes (9 to 0.05 μm; Ted Pella and Buehler) to obtain a mirror finish.

Fluorochrome-labeled surfaces were visualized using confocal laser scanning microscopy (Leica Stellaris DMI8, Leica Microsystems, Germany) with a 10× dry objective. Calcein was excited at 490 nm and detected at 500–568 nm; alizarin was excited at 579 nm and detected at 584–679 nm. Bone surfaces were imaged in reflectance mode to allow precise mapping of fluorochrome deposition. All images were acquired at 1024 × 1024 resolution, 400 Hz, with pinhole set to 1 Airy unit, and a single in-focus plane. Image contrast was adjusted uniformly using ImageJ (v1.54, NIH). Labeled surface areas were quantified on periosteal, endosteal, and total bone surfaces excluding any vasculature. All analyses were performed by investigators blinded to group assignment.

### 2.5 Raman spectroscopy

Matrix properties were assessed using Raman spectroscopy with a Horiba LabRam HR Evolution system (NIR, Japan) equipped with a 10× dry objective and 785 nm laser (100% power, model #: 0902B100-A, Horiba, Japan). Unembedded proximal femoral ends were thawed for four hours at 4°C prior to testing and maintained in a hydrated state using a custom holder that contains a tap water-saturated sponge. Spectra were acquired with an acquisition time of 5 s and 5 accumulations over a range of 300-1900 cm^−1^ at five equidistant points (250 μm apart) along the posterior shaft of the proximal femur, starting in line with the apex of the third trochanter. Data processing included 12th-order polynomial baseline correction in LabSpec6 (v6.7.2, Horiba France SAS, France). Spectral metrics included mineral-to-matrix ratio (MMR; ν_2_PO[area/Amide III area), carbonate-to-phosphate ratio (ν_1_CO[area/ν_1_PO[area), and crystallinity (inverse of FWHM of ν_1_PO[). Collagen maturity and secondary structure were assessed using Amide I sub-peak intensity ratios (I1670/I1640, I1670/I1610, I1670/I1690) determined by second-derivative analysis [6]. All Raman metrics were extracted using custom MATLAB code. Median values across five measurement sites were used per sample.

### 2.6 Serum biomarkers

Serum was analyzed for bone turnover markers C-terminal telopeptide of type I collagen (CTX-I; RatLaps® EIA, Immunodiagnostic Systems, UK) and procollagen type I N-terminal propeptide (P1NP; Rat/Mouse PINP EIA, Immunodiagnostic Systems, UK) using ELISA kits per manufacturer instructions. Absorbance was read using a microplate spectrophotometer (SpectraMax® iD3 instrument, SoftMax® Pro 7 7.1.2 software, Molecular Devices, USA).

### 2.7 Untargeted metabolomics

Untargeted metabolomics was performed to investigate mechanisms underlying the observed declines in cortical bone material properties. Metabolites were extracted according to the protocol described by Welhaven et al. [28]. Briefly, snap frozen proximal femoral halves were pulverized under liquid nitrogen using a mortar and pestle. Metabolites were extracted in 500uL of −20°C 3:1 HPLC grade methanol: acetone, followed by 5 cycles of 1-minute vortexing and 4-minute incubation at −80°C to facilitate macromolecule precipitation. Samples were then stored overnight at −80°C to ensure complete precipitation.

The following day, samples were centrifuged at 13,000 x g for 10 minutes at 4°C to remove cellular debris. Supernatant containing extracted metabolites was collected and dried via vacuum concentration. Metabolites were resuspended in 1:1 HPLC grade acetonitrile: water prior to analyses.

Mass spectrometry was performed in positive mode HILIC electrospray ionization liquid chromatography coupled MSE™ mass spectrometry. Data collection was performed on a Waters Synapt-XS Q-TOF with Waters I-Class UPLC unit and an Acquity Premier BEH Amide HILIC column. Scanning was performed for m/z values 50-1200 with a collision energy of 15-25eV. Data dependent analysis was run on a pooled sample for the top 4 peaks in each scan cycle excluding common false peaks with the same settings as MSE™. Resulting data were TIC normalized and tentatively identified (i.e., identified without standard confirmation) using Progenesis QI software (Waters Corporation, USA). Processed data were log transformed, z-scaled, and run through differential abundance testing, visualization (PCA, PLS-DA, hierarchical clustering), and pathway enrichment analysis pipeline using MetaboAnaylst 6.0 [29].

### 2.8 Statistical analyses

Statistical analyses were conducted in Minitab (v22, Minitab LLC, USA). Two-way ANOVAs were used to test the main effects of age, sex, and their interaction. Variables were assessed for normality and homogeneity of variance; data were Box-Cox transformed where appropriate. Grubbs’ test (α = 0.05) was used to identify and remove outliers. Parallel analyses without transformations or outlier removal were conducted for comparison. *Post hoc* tests were corrected for multiple comparisons using the Bonferroni method. Statistical significance was set at p < 0.05. Principal component analysis (PCA) and K-means clustering were carried out in a custom python code. Analysis included all measures of bone microstructure and tissue material quality (i.e., flexural testing, Raman spectroscopy, µCT). The value of k was chosen via the elbow method. Metabolite features were log transformed and z-scaled prior to analysis. Features were determined to be significant through the combination of a Fischer’s FDR-corrected exact t-test (α = 0.05) and |log_2_(Fold Change) ≥ 1|. Pathway enrichment utilized mummichog MSEA (α = 0.25). *Post hoc* power analyses were performed using G*Power 3.1.9.7 (Heinrich Heine University, Germany) to estimate effect sizes in bone outcomes (Cohen’s f, **ST1**) [30]. Metabolite abundance and bone outcomes for correlation analysis were from the same individuals.

## 3.0 Results

### 3.1 Males show greater age-related changes to bone geometry and microstructure than females

Both sexes exhibited cross-section expansion and age-related changes in cortical microarchitectural, although several outcomes showed clear sex-specific trajectories (**Table 1**, **Figure 1**). Aging induced significant alterations in mid-diaphyseal cortical architecture, with multiple parameters demonstrating age-by-sex interactions. Age and sex had interactive effects on pMOI (p = 0.042) and I_min_ (p = 0.006), with males exhibiting greater age-related changes in cortical geometry. From 7 to 22 months of age, females showed a 22.5% increase in pMOI and no change in I_min_, whereas males exhibited a 30.6% increase in pMOI and a 34.2% increase in I_min_.

**Figure 1.**
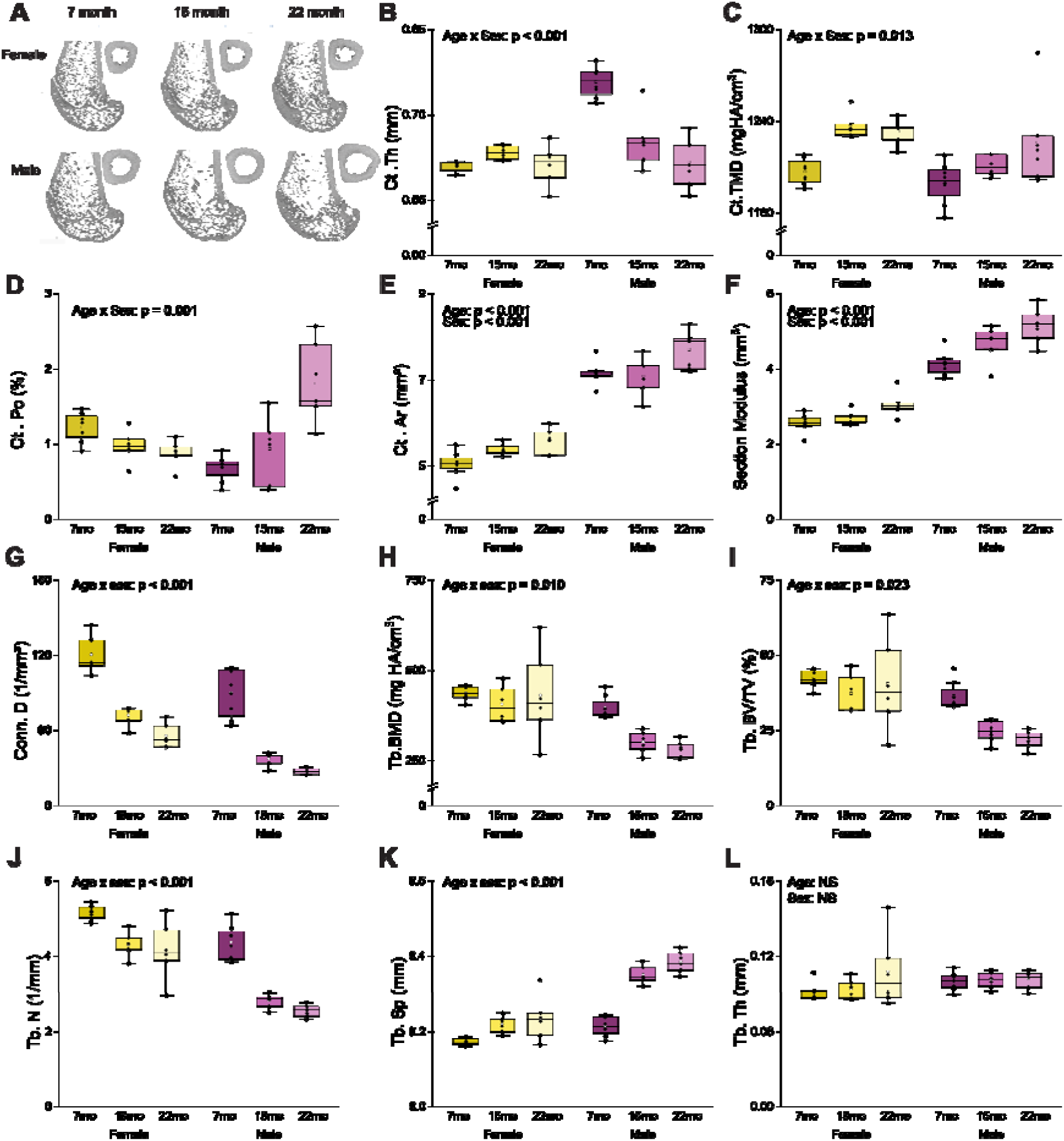
Age-related changes in bone microstructure and cortical geometry differed between males and females. (A) Representative µCT images for each group. Cortical bone outcomes include (B) cortical thickness (Ct. Th), (C) cortical tissue mineral density (Ct. TMD), (D) cortical porosity (Ct. Po), (E) cortical area (Ct. Ar), and (F) section modulus (I_min_/c_min_). Trabecular bone properties outcomes include (G) connectivity density (Conn. D), (H) trabecular bone mineral density (Tb. BMD), (I) trabecular bone volume fraction (Tb. BV/TV), (J) trabecular number (Tb. N), (K) trabecular separation (Tb. Sp)., and (L) trabecular thickness (Tb. Th). P-values from two-way ANOVA are reported for variables exhibiting significant main effects or interactions. Boxes represent the interquartile range (IQR), while whiskers represent the data range within 1.5 IQR. The solid line represents the median group value, and the transparent dot represents the group’s mean. Samples per group range from 6 to 8.

**Table 1.**
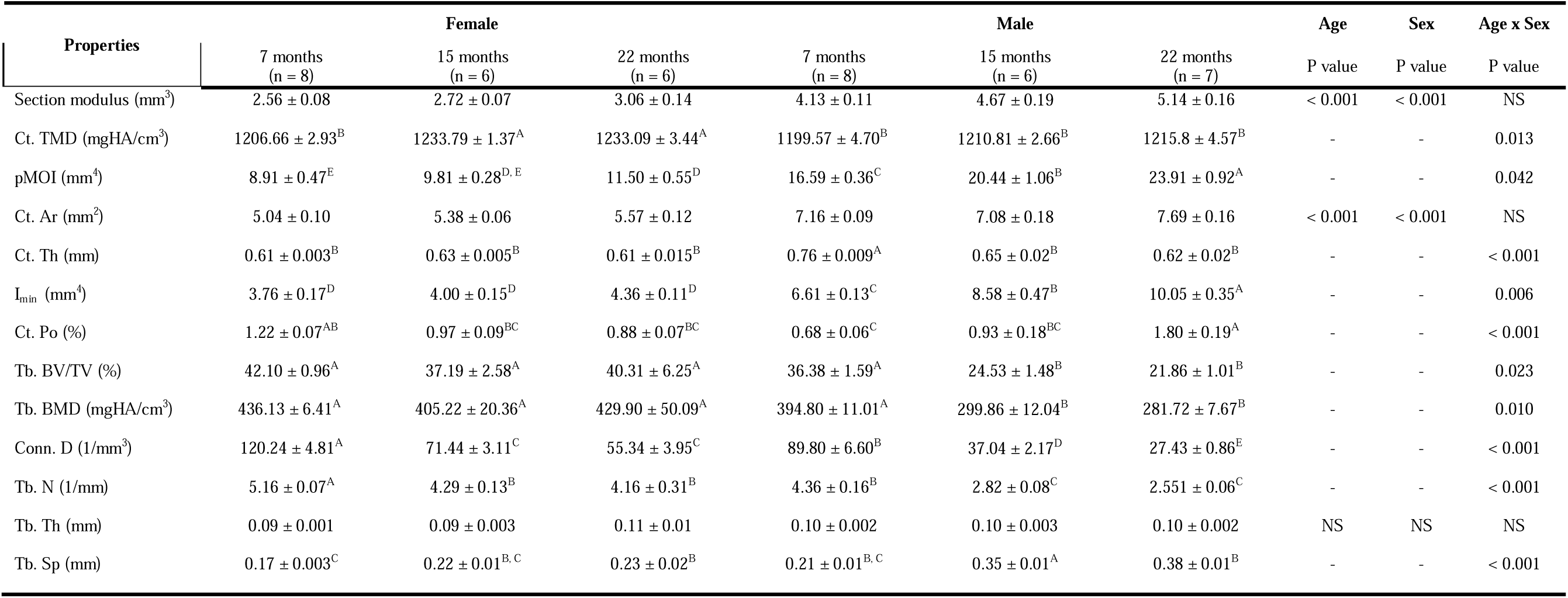
Bone microstructure and cortical geometry assessed by micro-tomography. P-values correspond to two-way ANOVA results. Means that do not share a common letter differ significantly for age-by-sex interactions. Data are presented as mean ± standard error. Non-significant results are denoted as NS unless the p-value is ≤ 0.10.

| Properties | Female |  |  | Male |  |  | Age | Sex | Age x Sex |
| --- | --- | --- | --- | --- | --- | --- | --- | --- | --- |
|  | 7 months<br>(n = 8) | 15 months<br>(n = 6) | 22 months<br>(n = 6) | 7 months<br>(n = 8) | 15 months<br>(n = 6) | 22 months<br>(n = 7) | P value | P value | P value |
| Section modulus (mm<sup>3</sup>) | 2.56 ± 0.08 | 2.72 ± 0.07 | 3.06 ± 0.14 | 4.13 ± 0.11 | 4.67 ± 0.19 | 5.14 ± 0.16 | < 0.001 | < 0.001 | NS |
| Ct. TMD (mgHA/cm<sup>3</sup>) | 1206.66 ± 2.93<sup>B</sup> | 1233.79 ± 1.37<sup>A</sup> | 1233.09 ± 3.44<sup>A</sup> | 1199.57 ± 4.70<sup>B</sup> | 1210.81 ± 2.66<sup>B</sup> | 1215.8 ± 4.57<sup>B</sup> | - | - | 0.013 |
| pMOI (mm<sup>4</sup>) | 8.91 ± 0.47<sup>E</sup> | 9.81 ± 0.28<sup>D,E</sup> | 11.50 ± 0.55<sup>D</sup> | 16.59 ± 0.36<sup>C</sup> | 20.44 ± 1.06<sup>B</sup> | 23.91 ± 0.92<sup>A</sup> | - | - | 0.042 |
| Ct. Ar (mm<sup>2</sup>) | 5.04 ± 0.10 | 5.38 ± 0.06 | 5.57 ± 0.12 | 7.16 ± 0.09 | 7.08 ± 0.18 | 7.69 ± 0.16 | < 0.001 | < 0.001 | NS |
| Ct. Th (mm) | 0.61 ± 0.003<sup>B</sup> | 0.63 ± 0.005<sup>B</sup> | 0.61 ± 0.015<sup>B</sup> | 0.76 ± 0.009<sup>A</sup> | 0.65 ± 0.02<sup>B</sup> | 0.62 ± 0.02<sup>B</sup> | - | - | < 0.001 |
| I<sub>min</sub> (mm<sup>4</sup>) | 3.76 ± 0.17<sup>D</sup> | 4.00 ± 0.15<sup>D</sup> | 4.36 ± 0.11<sup>D</sup> | 6.61 ± 0.13<sup>C</sup> | 8.58 ± 0.47<sup>B</sup> | 10.05 ± 0.35<sup>A</sup> | - | - | 0.006 |
| Ct. Po (%) | 1.22 ± 0.07<sup>AB</sup> | 0.97 ± 0.09<sup>BC</sup> | 0.88 ± 0.07<sup>BC</sup> | 0.68 ± 0.06<sup>C</sup> | 0.93 ± 0.18<sup>BC</sup> | 1.80 ± 0.19<sup>A</sup> | - | - | < 0.001 |
| Tb. BV/TV (%) | 42.10 ± 0.96<sup>A</sup> | 37.19 ± 2.58<sup>A</sup> | 40.31 ± 6.25<sup>A</sup> | 36.38 ± 1.59<sup>A</sup> | 24.53 ± 1.48<sup>B</sup> | 21.86 ± 1.01<sup>B</sup> | - | - | 0.023 |
| Tb. BMD (mgHA/cm<sup>3</sup>) | 436.13 ± 6.41<sup>A</sup> | 405.22 ± 20.36<sup>A</sup> | 429.90 ± 50.09<sup>A</sup> | 394.80 ± 11.01<sup>A</sup> | 299.86 ± 12.04<sup>B</sup> | 281.72 ± 7.67<sup>B</sup> | - | - | 0.010 |
| Conn. D (1/mm<sup>3</sup>) | 120.24 ± 4.81<sup>A</sup> | 71.44 ± 3.11<sup>C</sup> | 55.34 ± 3.95<sup>C</sup> | 89.80 ± 6.60<sup>B</sup> | 37.04 ± 2.17<sup>D</sup> | 27.43 ± 0.86<sup>E</sup> | - | - | < 0.001 |
| Tb. N (1/mm) | 5.16 ± 0.07<sup>A</sup> | 4.29 ± 0.13<sup>B</sup> | 4.16 ± 0.31<sup>B</sup> | 4.36 ± 0.16<sup>B</sup> | 2.82 ± 0.08<sup>C</sup> | 2.551 ± 0.06<sup>C</sup> | - | - | < 0.001 |
| Tb. Th (mm) | 0.09 ± 0.001 | 0.09 ± 0.003 | 0.11 ± 0.01 | 0.10 ± 0.002 | 0.10 ± 0.003 | 0.10 ± 0.002 | NS | NS | NS |
| Tb. Sp (mm) | 0.17 ± 0.003<sup>C</sup> | 0.22 ± 0.01<sup>B,C</sup> | 0.23 ± 0.02<sup>B</sup> | 0.21 ± 0.01<sup>B,C</sup> | 0.35 ± 0.01<sup>A</sup> | 0.38 ± 0.01<sup>B</sup> | - | - | < 0.001 |

Age and sex interactions were also observed for Ct. Th (p < 0.001), Ct. Po (p < 0.001), and Ct. TMD (p = 0.013). In females, Ct. Th and Ct. Po remained unchanged, while Ct. TMD increased modestly by 2.1% from 7- to 22-mo. In contrast, males exhibited an 18.6% decrease in Ct. Th and a 164.7% increase in Ct. Po, with no change in Ct. TMD over the same interval. Aging was associated with cortical expansion, reflected by significant increases in Ct. Ar (p < 0.001) and section modulus (p < 0.001). Compared with 7-mo rats, 22-mo rats exhibited a 9.15% increase in Ct. Ar and a 19.9% increase in section modulus. Sex exerted an additive effect, with males exhibiting greater Ct. Ar (p < 0.001) and section modulus (p < 0.001) across ages.

Aging was also associated with marked deterioration of trabecular microarchitecture, with more pronounced effects in males. Significant interactions between age and sex were found for Tb. BV/TV (p = 0.023), Conn. D. (p < 0.001), Tb. N. (p = 0.021), Tb. BMD (p = 0.010), and Tb. Sp (p < 0.001). From 7 to 22 months, males exhibited a 39.9% reduction in Tb. BV/TV, whereas females showed no age-related change. Over the same interval, females experienced a 50.4% decrease in Conn. D., a 19.4% decrease in Tb. N, no change in Tb. BMD, and a 26.1% increase in Tb. Sp. In contrast, males exhibited a 69.5% decrease in Conn. D., a 41.4% decrease in Tb. N., a 28.6% decrease in Tb. BMD, and a 44.0% increase in Tb. Sp.

For several cortical and trabecular measures, including Section Modulus, Ct. TMD, pMOI, Ct. Th, I_min_, Tb. BV/TV, Tb. BMD, Conn. D, Tb. N, and Tb. Sp, age-related changes were already evident by 15 months compared with 7 months. In contrast, changes in Ct. Ar and Ct. Po were not apparent until 22 months. Full statistical results are available in **Table 1**.

### 3.2 Both sexes exhibit reduced fracture resistance by midlife, with greater declines in males

Whole-bone flexural properties of the femur declined by early midlife in a sex-dependent manner (**Table 2**, **Figure 2**). Most material properties demonstrated additive effects of age and sex, whereas modulus exhibited a significant age x sex interaction (p = 0.002). Females maintained modulus across age, while males exhibited a 24.45% decrease from 7-mo to 22-mo.

**Figure 2.**
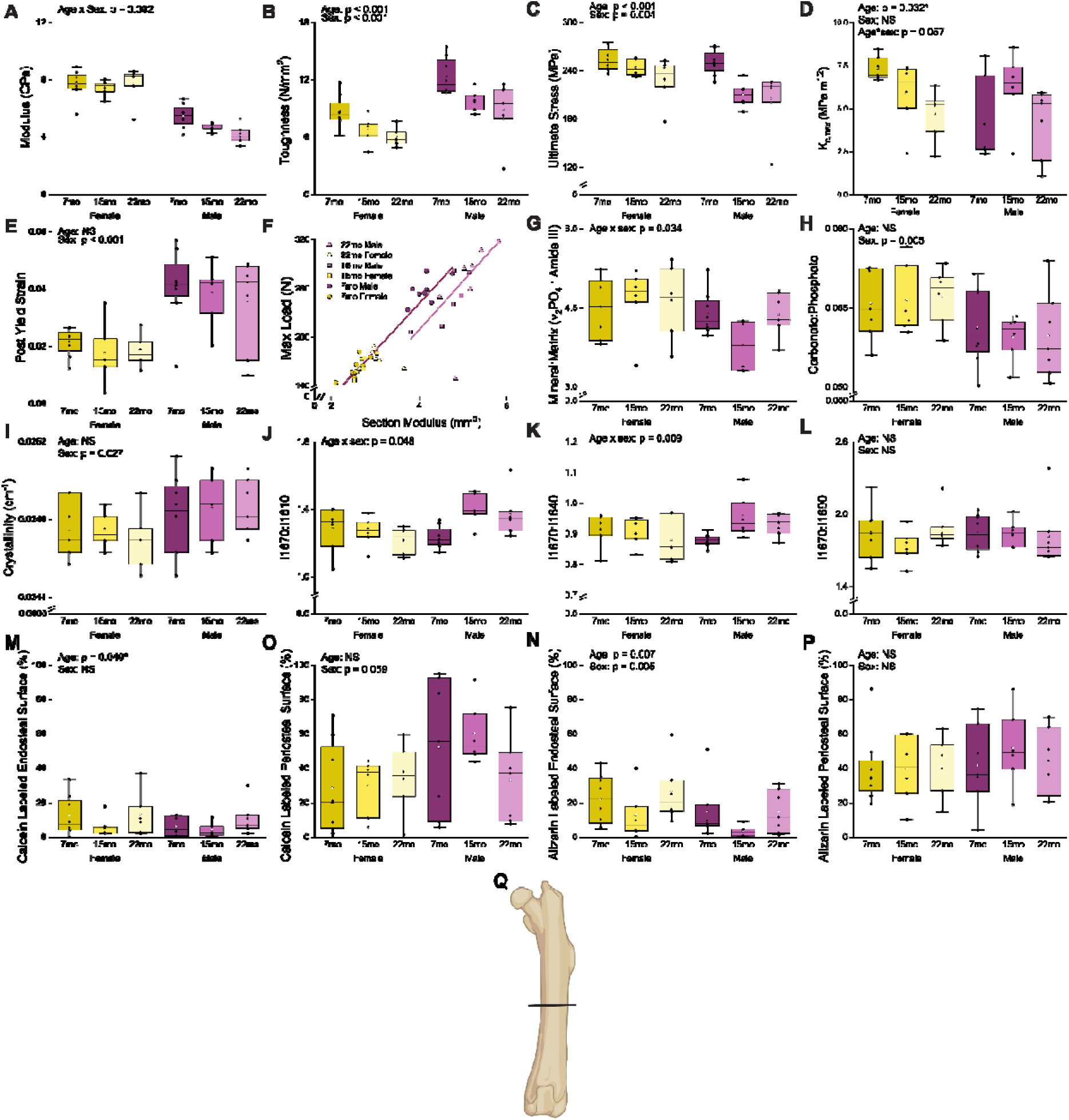
Bone material properties exhibit age-related and sex-specific differences. Flexural testing outcomes include (A) modulus, (B) toughness, (C) ultimate stress, (D) K_c,max_, (E) post yield strain, and (F) geometry scaled strength, assessed by peak load versus section modulus. Raman derived material properties were (G) mineral-to-matrix ratio (ν_2_PO□area/Amide III area), (H) carbonate: phosphate ratio (ν_1_CO_3_ area/ ν_1_PO_4_ area), (I) crystallinity (1/FWHM ν_1_PO_4_), Amide I subpeak ratios (J) I1670:I1610, (K) I1670:I1640, and (L) I1670:I1690. Dynamic histomorphometry was assessed for (M) the endosteal surface 10 days and (N) 3 days prior to euthanasia as well as (O) the periosteal surface 10 days and (P) 3 days prior to euthanasia. (Q) Location of histomorphometry analysis. Representative stitched images, total surface labeling, and the serum CTX1:P1NP ratio are shown in **Figure S2**. For plots A-E and G-P, p-values from two-way ANOVA are reported for variables exhibiting significant main effects or interactions. Boxes represent the IQR, while whickers represent the data range within 1.5 IQR. The solid line represents the median group value, and the transparent dot represents the group’s mean. Samples per group range from 3 to 8. *Indicates effects that were no longer statistically significant following family-wise error correction.

**Table 2.**
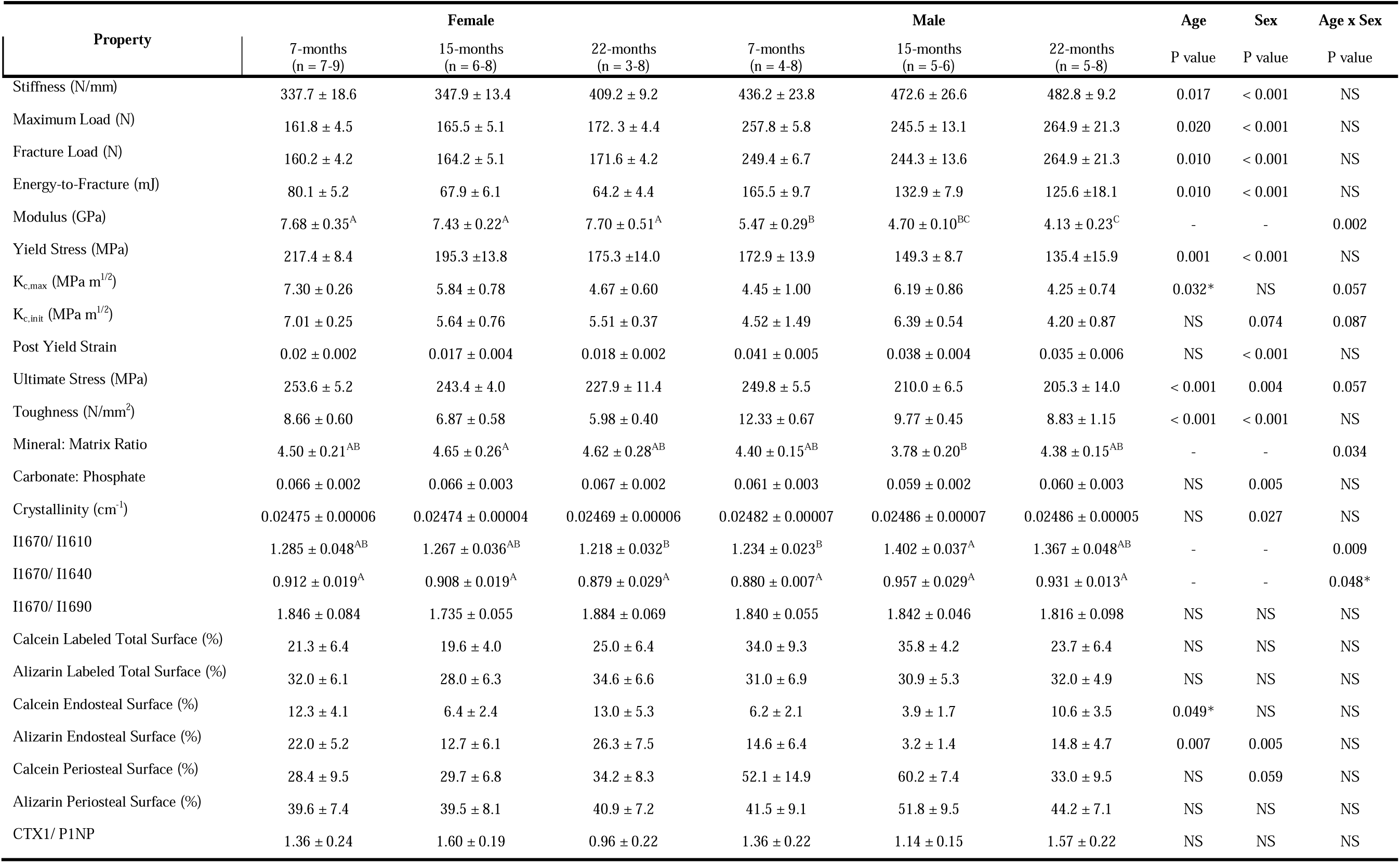
Material properties derived from three-point bending. Matrix property ratios extracted from Raman spectroscopy. Bone turnover and activity assessed by serum biomarkers and quantitative histomorphometry. P-values correspond to two-way ANOVA results. Means that do not share a common letter differ significantly for age-by-sex interactions. Data are presented as mean ± standard error. Non-significant results are denoted as NS unless the p-value is ≤ 0.10. Load-displacement curves, stress-strain curves, K_c,init_ boxplots, and a representative image for fracture toughness testing are provided in supplemental figures **S3**-**S6**, respectively. *Indicates effects that were no longer statistically significant following family-wise error correction.

Ultimate stress (age: p < 0.001, sex: p = 0.004) and yield stress (age: p = 0.001, sex: p < 0.001) declined with age and differed between sexes as main effects, with no significant interaction. Males regardless of age exhibited 8.60% lower ultimate stress and 28.97% lower yield stress compared with females. Aging reduced ultimate stress and yield stress by 14.30% and 21.17%, respectively, when comparing 22-mo to 7-mo rats. Following outlier removal and transformation to meet ANOVA assumptions, ultimate stress showed a trend toward an age x sex interaction (p = 0.057), with males exhibiting consistently lower values by 15-mo and females declining progressively through 22-mo; however, this interaction did not meet the prespecified *a priori* threshold for significance and may reflect limited statistical power (**ST1**).

Bone toughness declined progressively with age (p < 0.001), decreasing 28.39% from 7-mo to 22-mo. A significant main effect of sex was also observed (p < 0.001), with males exhibiting 29.88% greater toughness than females. Post-yield strain was influenced only by sex (p < 0.001), with males demonstrating 100.34% greater post-yield strain than females. Strain hardening was observed during notch fracture testing in eight individuals (7-mo M: 2, 15-mo M: 1, 22-mo F: 3, 22-mo M: 2; see **S1**). These individuals were excluded from K_c,init_ analysis. K_c,init_ may have an effect of sex (p = 0.074) and an interaction of age and sex (p = 0.087) but did not reach significance. Statistical analysis has the potential to be underpowered, especially since several data points were excluded (n = 3 – 7 per group, **ST1**). K_c,max_ decreased with age (p = 0.032) but did not meet significance criteria following family-wise correction. K_c,max_ may be affected by the interaction of age and sex (p = 0.057). Females may have lower K_c,init_ and K_c,max_ with age, while males do not show differences in these measures with age.

To evaluate strength differences between groups after accounting for femur cross-sectional size differences, peak load was regressed against section modulus. All female age groups and 7-mo males exhibited a similar linear relationship between bone size and strength (i.e., all on same line). In contrast, both 15-mo and 22-mo males were on a different, lower line. These data show that bone strength is reduced in aged males compared with other groups and not because of size differences in the bones (**Figure 2F**). Full statistical results are available in **Table 2**.

### 3.3 Females maintain tissue-scale matrix properties through 22-mo, while males exhibit transient midlife decline in collagen integrity

Raman spectroscopy revealed age- and sex-dependent differences in cortical bone matrix properties of the femur (**Table 2**, **Figure 2**). Age and sex had interactive effects (p = 0.034) on the mineral-to-matrix ratio (MMR, ν_2_PO[area/Amide III area). Males had an 18.7% lower MMR compared with age-matched female at 15-mo with no sex differences observed at 7-mo or 22-mo.

Indices of mineral maturity differed by sex but were not influenced by age. The carbonate: phosphate ratio (ν_1_CO_3_ area/ ν_1_PO_4_ area; p = 0.005) was 9.4% lower in males than females, while crystallinity (1/FWHM ν_1_PO_4_; p = 0.027), was 0.4% higher in males.

Collagen-related spectral features demonstrated aging trajectories that differed by sex. Age and sex had interactive effects on amide I subpeak intensity ratios I1670:I1610 and I1670:I1640 (p = 0.048 and p = 0.009, respectively). Females maintained stable ratios across age, whereas males exhibited a transient increase in both measures at 15-mo compared with 7-mo. *Post hoc* testing did not show differences between individual contrasts for I1670:I1640, following familywise error correction. Relative to 7-mo males, I1670:I1610 increased by 13.6% and I1670:I1640 increased by 8.8% at 15-mo, while females exhibited no changes in either ratio with age. No age- or sex-related differences were detected for the I1670:I1690 ratio (p > 0.10). Full statistical results are available in **Table 2**.

### 3.4 Age and sex influence local, but not serum, measures of bone turnover

Dynamic histomorphometry of fluorochrome-labeled surfaces revealed site-specific and time-dependent differences in cortical turnover. (**Table 2**, **Figure 2**). When periosteal and endocortical sites were analyzed together, mineralizing surface did not differ by age or sex for either calcein or alizarin labeling (p > 0.010 for both). In contrast, endosteal turnover demonstrated both age- and sex-related effects. Endosteal alizarin-labeled surface was influenced by age (p = 0.007) and sex (p = 0.005), while calcein labeling showed an effect of age (p = 0.049). However, following correction for family-wise error, age-related differences in calcein-labeled endosteal surface were no longer statistically significant.

Relative to 7-mo rats, 15-mo animals exhibited the lowest endosteal activity, with a 57.3% reduction in alizarin labeling and a 45.3% reduction in calcein labeling. Females exhibited 82.0% greater endosteal activity than males at the later labeling point, three days prior to euthanasia, whereas no sex differences were observed at the earlier labeling point. Periosteal activity remained stable across age groups (p > 0.10 for both labels). Calcein-labeled periosteal surface trended toward higher values in males (56.7% greater than females), although this difference did not reach statistical significance (p = 0.059). Alizarin-labeled periosteal surface did not differ by sex (p > 0.10).

Systemic markers of bone turnover, serum CTX-I (resorption) and P1NP (formation), did not differ by age or sex, and the P1NP:CTX-I ratio remained similar among groups (p > 0.10), indicating no detectable differences in systemic turnover across these ages. Bone turnover measures exhibited weak associations with µCT, flexural testing, and Raman spectroscopy results (|Pearson’s r| < 0.2). Full statistical results are available in **Table 2**

### 3.5 Principal components analysis reveals age- and sex-specific clustering of bone quality phenotypes

To identify the primary contributors to group-level differences, principal component analysis was performed using z-scaled bone quality variables. This analysis revealed three distinct clusters (**S7**): one comprising all female samples regardless of age, one comprising 7-mo males, and one comprising 15- and 22-mo males. Sample identities were mapped unambiguously to these clusters.

The first principal component (PC1) explained 50.5% of the total variance and was dominated by variables related to bone size and cortical geometry, including Tt. Ar, I_min_, pMOI, section modulus, and I_max_ (see **ST2** for full variable weighting). These variables produced a clear separation by sex, with larger bones plotted on the right side of **S7**.

The second principal component (PC2) explained an additional 12.5% of the variance and was driven primarily by measures related to fracture resistance, including toughness, Ct. Th, energy at fracture, ultimate stress, and Ct. Ar/Tt. Ar. PC2 separated young and midlife males, reflecting an age-associated decline in fracture resistance within males. In contrast, female samples did not change along this component. Collectively, these results indicate that aging more strongly influences multiple dimensions of bone quality (e.g., cortical geometry and material properties) in males than in females by midlife in this strain.

### 3.6 The cortical metabolome of males is more affected by age than that of females

Untargeted metabolomics of the contralateral femur revealed that age-related metabolic changes were more pronounced in males than females, with sex differences increasing across age. (**Figure 3**). No age-related main effects were found but age-related effects were present in a sex-specific manner. In both sexes, no age-related differences were detected at 15-mo of age compared to 7-mo. By 22-mo, males exhibited metabolic remodeling, with 73 metabolites upregulated and two downregulated compared to 7-mo males. These changes were associated with enrichment of amino acid metabolism (cysteine, methionine, arginine, lysine) and nucleotide metabolism (pyrimidine, purine, amino sugar, nucleotide sugar). In contrast, females exhibited only one significantly upregulated metabolite over the same period, with no enriched pathways identified.

**Figure 3.**
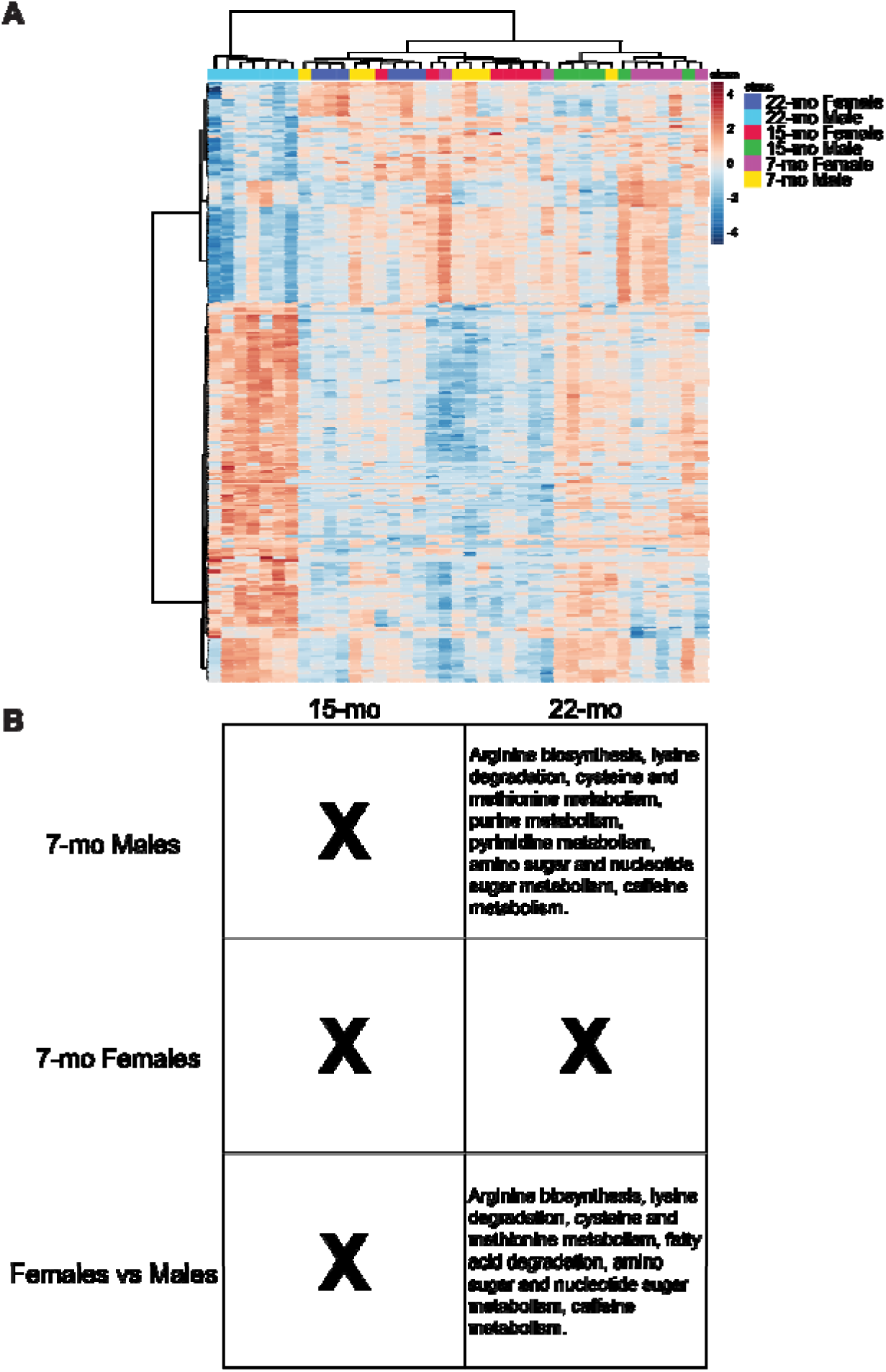
Cortical metabolome. (A) Heatmap of the top 250 metabolite features ranked by p-value with Ward clustering of both samples and features. Warmer colors represent greater abundance, while cooler colors represent less abundance. (B) Tables of KEGG pathways enriched from significant features due to age-related effects in males and females alongside sex differences in age. Volcano plots for significant comparisons; all-feature heatmap, PCA, and PLS-DA plots are in **S8**.

Sex differences in metabolite abundance were increased with age. Regardless of age, 29 metabolite features differed by sex, with 28 higher in males and 1 higher in females; however, these differences did not result in enriched pathways. Age-matched comparisons revealed increasing divergence between sexes. At 7-mo, 15 metabolites differed between males and females, with one higher in females. At 15-mo, 21 metabolites differed between males and females, all of which were higher in males. At 22-mo, 116 features were significantly different between sexes, including 28 that were higher in females. At this time point, 22-mo males exhibited enrichment of amino acid metabolism (cysteine, methionine, arginine, lysine degradation), nucleotide metabolism (amino sugar and nucleotide sugar), and fatty acid degradation compared to age-matched females. No pathways were enriched in females compared with males at any age.

Correlation analyses of metabolite features and bone quality outcomes identified multiple moderate associations (e.g., Variables [Pearson’s r]: Ct. Ar/Tt. Ar-Niacinamide [–0.72], Ct. Po-Histidine [0.63], Modulus-Galabiose [-0.73], Ultimate Stress-Niacinamide [-0.60], Crystallinity-Galabiose [0.56]). Metabolites that are correlated with bone outcomes include niacinamide; sugars and advanced glycation end products; taurine; C16 phospholipids and carnitines; xanthine and hypoxanthine; and various amino acids like histidine. Full correlations results are available in the supplemental materials.

## 4.0 Discussion

The goal of this study was to evaluate whether the F344 × BN F1 rat represents a suitable translational model for investigating age-related deterioration of bone matrix and its contribution to declining fracture resistance. Using male and female rats spanning skeletal maturity to midlife, we demonstrate that tissue-scale material properties, including strength and toughness, decline by middle age in this strain, consistent with impaired fracture resistance. Although males and females exhibited comparable reductions in fracture resistance by later ages, the trajectories of microarchitecture, cortical geometry, and tissue-scale material properties changes differed between sexes. Importantly, bone strength accounting for bone size demonstrates clear differences in aging trajectory for male and female rats visualized by **Figure 2F**.

In males, the onset of reduced fracture resistance coincided with deterioration of both microarchitecture and matrix properties. These changes included cortical thinning, increased cortical porosity, trabecular degradation, reductions in mineral-to-matrix ratio, and transient alterations in collagen integrity indices during midlife [31]. Together, these findings suggest coordinated age-related changes in matrix properties and organization that contribute to declining fracture resistance (i.e., decreased strength and toughness). In contrast, females preserved bone mineral density, cortical geometry, and cortical microarchitecture across age yet still exhibited significant reductions in tissue-scale fracture resistance. These findings indicate that in females, declining fracture resistance is not driven by changes in cortical mineral content, assessed by µCT and Raman, or structure but instead reflects deterioration in matrix quality. Thus, male F344 × BN F1 rats exhibit a combined bone loss and matrix deterioration phenotype, whereas females primarily exhibit matrix-driven decline by 22-months-of-age. This divergence captures key features of skeletal aging relevant to translational investigation while occurring within a strain characterized by extended healthspan.

Cortical tissue metabolism also exhibited important sex-specific responses to aging. In males, aging was associated with enrichment of amino acid and nucleotide metabolism pathways, whereas females showed no pathway-level changes across age. Amino acid and nucleotide metabolism are central to bone cell function and may reflect alterations to multiple cellular processes including osteoclastogenesis, osteoblast proliferation, and collagen synthesis [32,33]. Sex differences in cortical bone metabolism increased with age, with pathway-level differences emerging at 22 months of age; when amino acid, nucleotide, and fatty acid metabolism are enriched in males with no corresponding enrichment in females. In addition to these pathways, *in vitro* and *in vivo* studies suggests that fatty acid metabolism, particularly β-oxidation, serves as an important energy source for bone formation and mineralization, which may contribute to the greater decline observed in aged males [34]. Correlation analysis further identified relationships between sugar abundance, taurine, citrate, and acylcarnitines with measures of bone material properties. Taurine has been shown to mitigate oxidative stress and modulate Wnt/β-catenin signaling in osteocytes, while citrate and acylcarnitines are linked to mitochondrial metabolism, suggesting mitochondrial function may serve an important role in maintenance of skeletal integrity with age [35]. This concept is supported by work in Cyclophilin D knockout mice, in which loss of function of the mitochondrial permeability transition pore attenuates age-related bone loss [36]. It remains unclear whether observed metabolic differences precede or follow declines in material properties, as feature selection was restricted to statistically significant metabolites.

The translational utility of this strain is further supported by its alignment with established features of skeletal aging observed across rodent models, while maintaining preserved systemic health later in life. The age-related declines in fracture resistance, tissue-scale mechanical properties, and matrix properties observed here parallel those reported in other rodent models including cortical thinning, trabecular bone loss, preservation of whole-bone flexural strength, reduced tissue-level strength, and declining toughness. However, the matrix deterioration observed in females in the absence of substantial bone loss represents a useful phenotype. This feature is particularly valuable for isolating the contribution of matrix quality to fracture resistance, independent of bone loss. The F344 x BN F1 rat maintains extended healthspan, enhancing its validity for translational aging research in contrast to many commonly used rodent models that develop alopecia, renal abnormalities, cardiovascular pathologies, and tumors earlier in life. As summarized in **Table 3** and **Figure 4**, this strain shares multiple skeletal aging features with both humans and other preclinical models while avoiding several confounding age-related pathologies. Additionally, variability in key material property outcomes, including ultimate stress, toughness, and elastic modulus, was comparable to or lower than that reported for other commonly used aged male rodent models [Ultimate stress CV: 18% F344xBNF1 (22-mo), 22% C57BL/6 (24-mo), 17% F344 (24-mo), 28% Wistar (24-mo); Toughness CV: 34% F344xBNF1 (22-mo), 92% C57BL/6 (24-mo), 34% F344 (24-mo); Elastic modulus CV: 16% F344xBNF1 (22-mo), 17% C57BL/6 (24-mo), 33% F344 (24-mo), 41% Wistar (24-mo)], which may reduce the number of animals required to achieve statistical power [7,10,15]. Together, these attributes position the F344 x BN F1 rat as a practical model for investigating matrix-driven skeletal aging.

**Figure 4.**
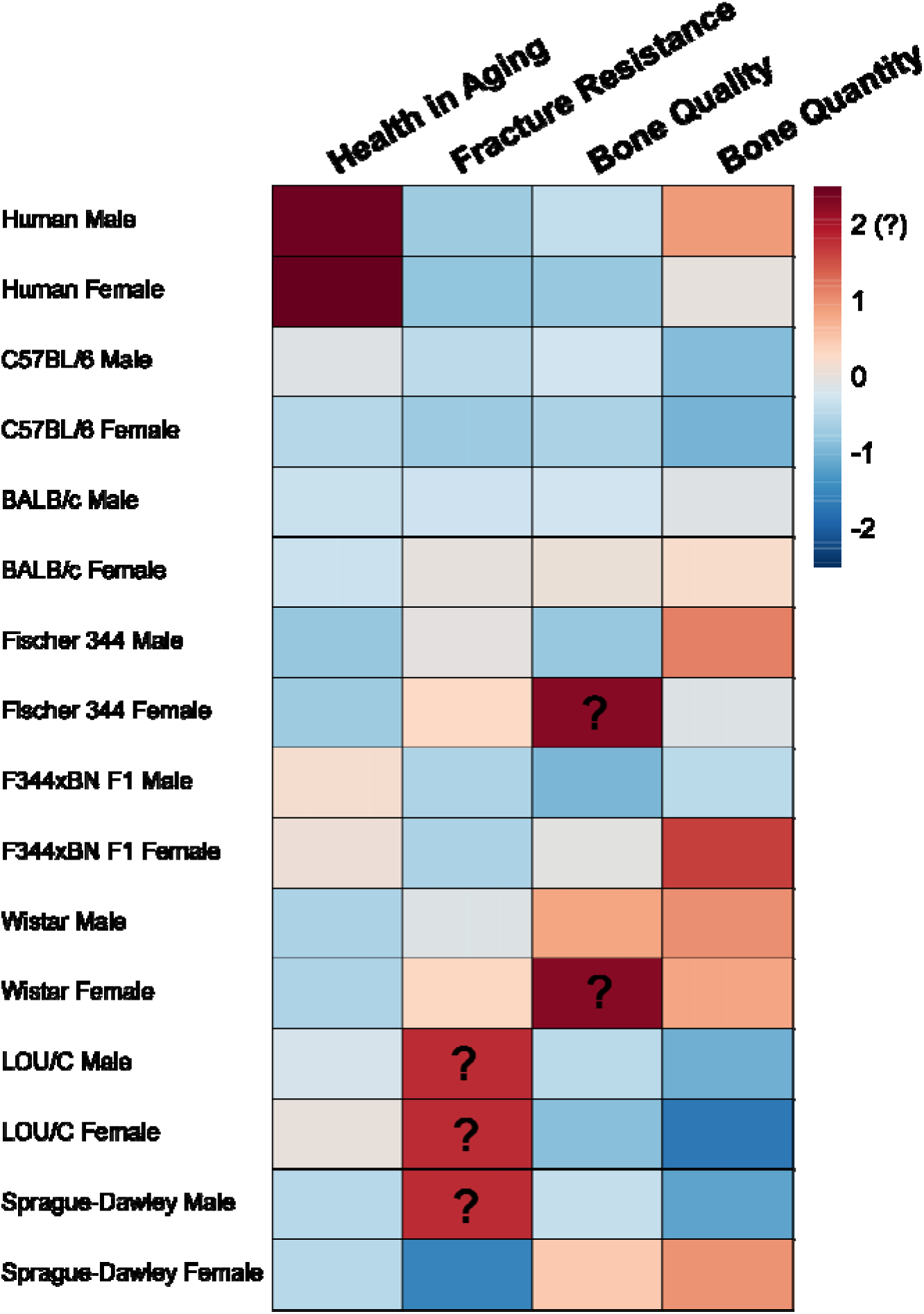
Comparison of skeletal aging features across rodent models and humans. Bone quantity, bone quality, fracture resistance, and overall health in aging represent major components of skeletal health relevant to translational research. The magnitude of age-related decline is illustrated using a heatmap, with cooler colors indicating greater decline in the given pseudovariable normalized to median lifespan. Positive values with a superimposed question mark, as shown for LOU/c fracture resistance, indicate unknown age-related changes. Pseudovariables represent z-scaled composites of age-related changes normalized to lifespan. Bone quantity is comprised of variables related to mineral density and percent bone area/volume; bone quality is comprised of variables related to microarchitecture and matrix properties; fracture resistance is comprised of variables related to strength, toughness, and yield; while health in aging is comprised of median lifespan and spontaneous tumorigenesis, with full component details provided in **Table ST3**. Femoral and other long-bone results were preferentially used in the preparation of this figure. Humans serve as the outgroup for skeletal and health decline in age. LOU/C and F344 x BN F1 rats show the greatest similarity to humans in healthspan-related decline but are still quite distinct from humans. C57BL/6 mice, LOU/c rats, and male SD rats exhibit the greatest decline in bone quantity across aging. F344 x BN F1 rats and C57BL/6 mice are the most similar to humans in fracture resistance, while various models exhibit a similar rate of age-related changes in bone quality to humans. Female F344 x BN F1 rats exhibit a pattern like human males of low age-related bone quantity loss and bone quality loss compared to the opposite sex but exhibit a similar loss of fracture resistance. Common features of skeletal aging include preservation of apparent strength through geometric remodeling, deterioration of trabecular microstructure, reduced tissue-scale fracture resistance, and alterations in matrix properties.

**Table 3.**
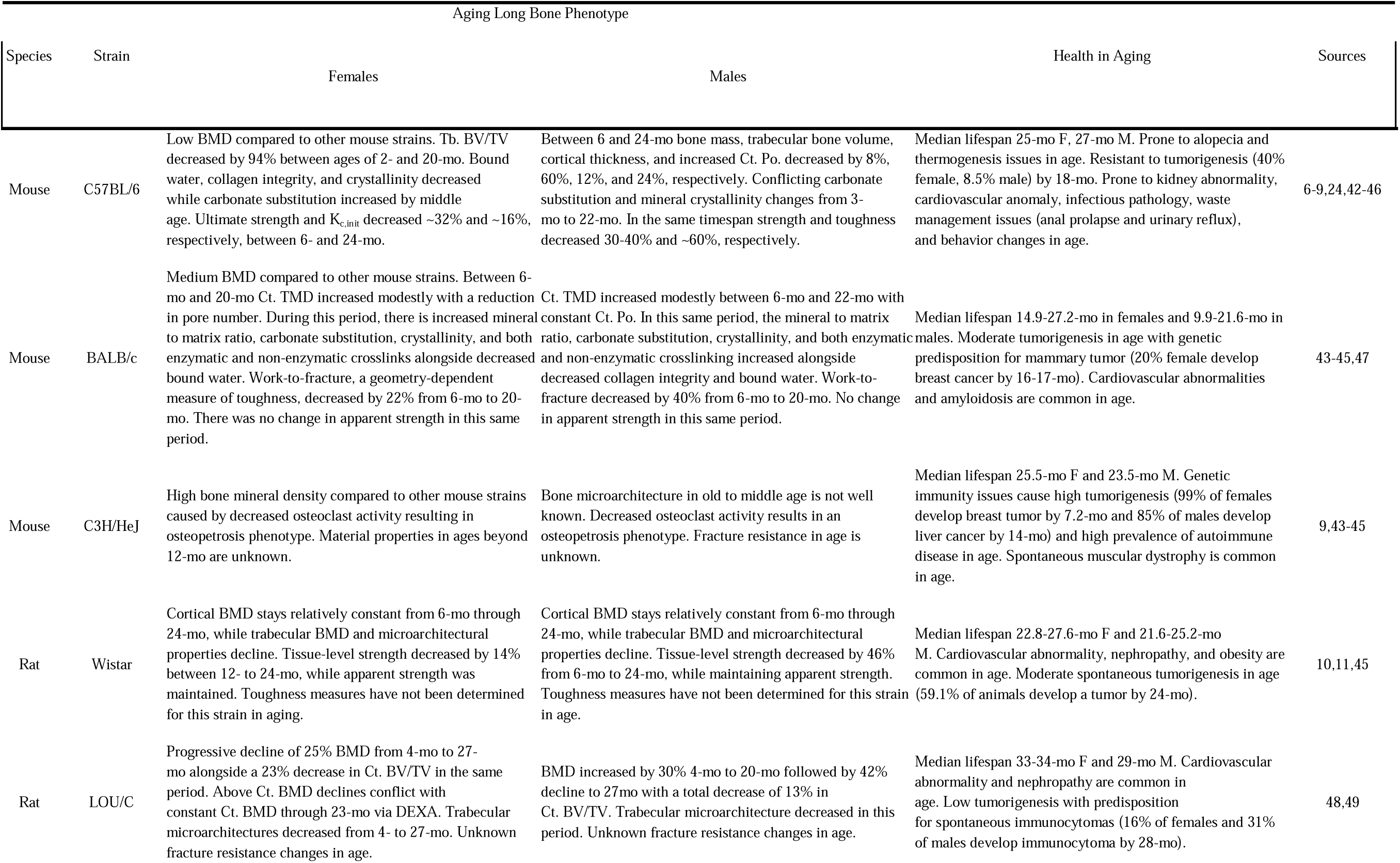
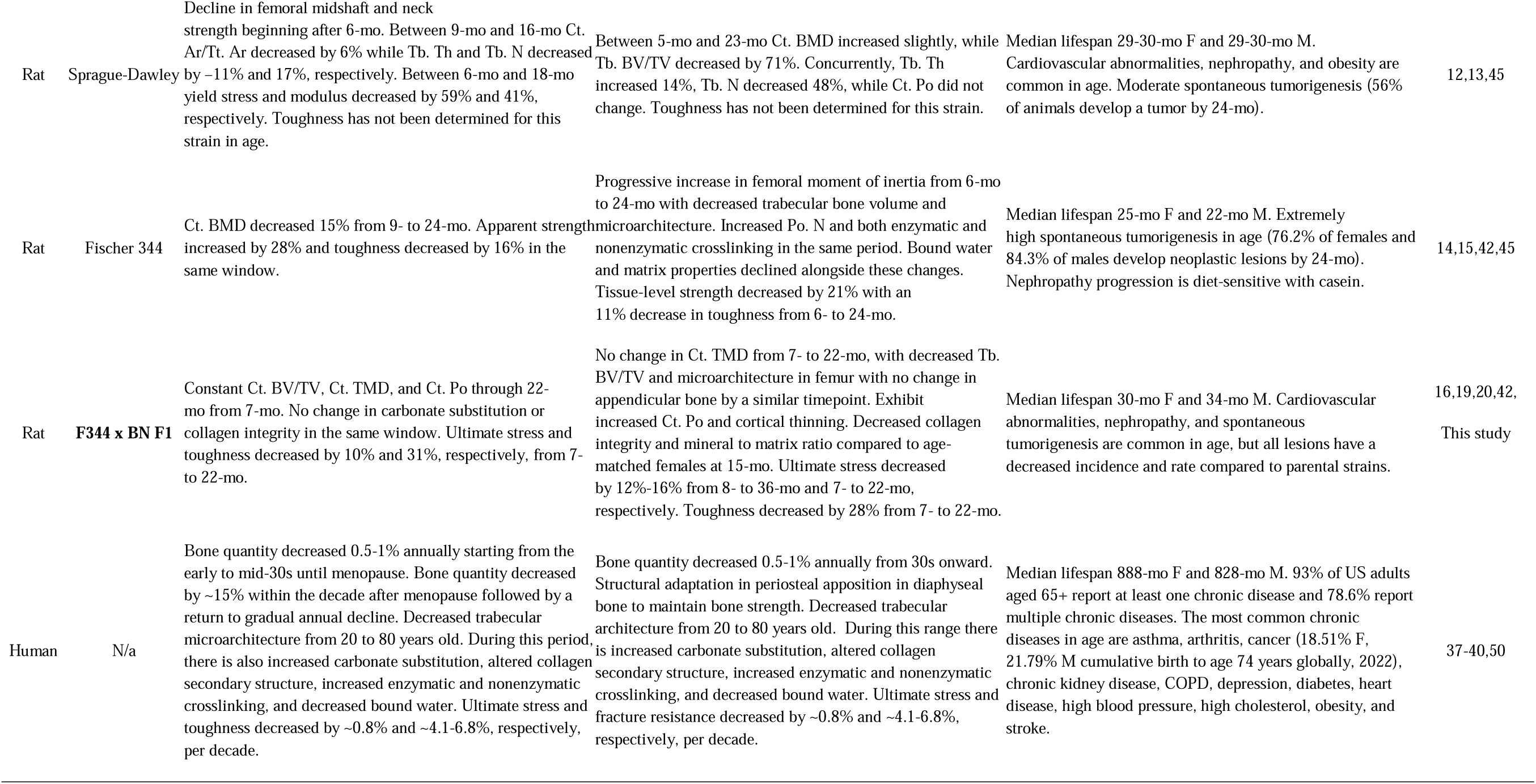
Summary of commonly used rodent models for aging and skeletal research along with human skeletal aging. Additional references are located within the supplementary document.

Sexual dimorphism in skeletal phenotype is well documented in both clinical and preclinical models and provides an opportunity to dissect the multi-factorial contributors to bone phenotype. Although the degree and nature of sexual dimorphism in this strain differ from humans in important aspects, both sexes offer complementary and informative models of skeletal aging. The female phenotype, characterized by declining fracture resistance despite preserved bone mass and microarchitecture, offers a unique platform for isolating the contribution of matrix quality to fracture resistance. This is particularly relevant given the growing recognition that bone mineral density alone is an incomplete predictor of fracture risk [2]. In this context, the present findings align with emerging work exploring matrix targeted therapeutic strategies for mitigating bone fragility in aging [4]. In contrast, males exhibit concurrent declines in bone mass, matrix quality, and material properties, a pattern that more closely resembles skeletal aging observed in humans and in several commonly used rodent models [37–40,7,10–15]. As such, male F344 × BN F1 rats provide a model well suited for studying the combined effects of bone loss, microarchitectural degradation, and matrix-level changes on fracture resistance, as well as for benchmarking matrix-focused interventions against more traditional osteopenic phenotypes.

Comparison of this strain’s skeletal aging phenotype with that of humans further supports its translational relevance, with the notable exception of postmenopausal osteoporosis. Shared features between F344 x BN F1 rats and humans include age-related increases in section modulus, declines in trabecular microarchitecture, reduction in collagen integrity, and decreased fracture resistance [37–40]. Differences between this model and human skeletal aging reflect limitations common to rodent models, including the absence of menopause and osteonal remodeling. Nevertheless, the combination of sex-specific aging trajectories and conserved features of bone aging across species allow F344 x BN F1 rats to address several persistent gaps in the skeletal aging literature. These include understanding why bone mass poorly predicts fracture risk in aging populations, elucidating how matrix deterioration contributes to mechanical fragility in the absence of substantial bone loss, and informing the development of matrix-targeting therapeutics.

This study has several limitations that should be considered. First, the assessment of bone matrix was not exhaustive. Important contributors to matrix quality including bound water content, enzymatic and non-enzymatic crosslinking, and perilacunar turnover were not evaluated and warrant investigation. Potential molecular mediators of matrix deterioration, such as oxidative stress, hormonal alterations, alterations in circulating factors, and changes in bone cell activity, were also not assessed and represent important targets for future mechanistic studies. In addition, female rats in this strain, as in other rat models, did not exhibit age-related bone loss; therefore, ovariectomy would be required to model estrogen-deficient osteoporosis, a condition not examined here [40]. The focus on middle age in this work was intentional, as this period is particularly relevant for identifying translationally actionable targets, but inclusion of later ages would be necessary to fully characterize skeletal aging into senescence. Finally, material properties derived from flexural testing rely on beam theory assumptions and are sensitive to differences in bone size and geometry, which are pronounced in this strain [41]. These estimates should therefore be interpreted with appropriate caution, particularly for sex-based comparisons.

In summary, this study demonstrates that F344 × BN F1 rats exhibit impaired bone fracture resistance by middle age, but with useful differences between females and males in aging. Over the period studied, males showed bone loss and matrix deterioration while females showed matrix deterioration without substantial bone loss. These divergent trajectories enable deconvolution of bone mass and bone quality contributions to fracture resistance and directly address a key limitation of many existing preclinical models. Collectively, the structural, mechanical, and matrix properties changes identified here support the F344 × BN F1 rat as a robust model for studying matrix-driven skeletal aging and for evaluating interventions aimed at preserving bone quality and fracture resistance across the healthspan.

## 5.0 Acknowledgments

This work is made possible by funding from the National Institute of Health (NIH) – Institute on Aging (SM, R21AG075402), NIH – National Institute of General Medical Sciences (P30GM154593), and the National Science Foundation (NSF) (CH, Career Award NSF 2340823). This work used the Animal Resources Center (RRID:SCR_026351), the Center for Biofilm Engineering Imaging Facility (RRID:SCR_009943), and Montana State Mass Spectrometry Facility (RRID: SCR_012482) which are supported by the Montana State University Office of Research and Economic Development, the NSF MRI Program (2018562), the M.J. Murdock Charitable Trust (202016116), the NIH – National Institute of General Medical Sciences (P20GM103474, S10OD28650), and the U.S. Department of Defense (77369LSRIP). We thank the staff of Montana State University’s Animal Resource Center, especially Tamara Marcotte and James Fox, for their assistance in providing excellent animal care. We also thank Donald Smith and Jesse Thomas for their work in ensuring the collection of high-quality mass spectrometry data. The content is solely the responsibility of the authors and does not necessarily represent the official views of funding agencies. Apologies to all the authors that could not be cited due to reference limitations.

## 6.0 Author Contributions

HH: software, investigation, formal analysis, data curation, writing—original draft, writing—review and editing, visualization; EK: investigation, formal analysis, writing—original draft, writing—review and editing; KB: methodology, software, investigation, formal analysis, writing—review and editing; SS: formal analysis, writing—original draft, writing—review and editing; CMH: software, conceptualization, formal analysis, resources, writing—original draft, writing—review and editing, funding acquisition; SAM: conceptualization, investigation, formal analysis, resources, writing—original draft, writing—review and editing, funding acquisition.

## 8.0 Data and Code Availability

The data and codes used may be requested from the corresponding author upon reasonable request.

## 9.0 Declarations of Generative AI

During the preparation of this work, the authors did not use generative AI.

## Data Availability

The data from this study are available from the corresponding author upon reasonable request.

## Ethics approval statement

The Montana State University Institutional Animal Care and Use Committee approved all animal procedures.

