## Supplementary material for "Bone microarchitecture and material properties decline differently across midlife for male and female F344 × BN F1 rats": ST1

**Supplementary Materials:**

ST1. Power Analysis Table for F344 x BN F1 Rat Bone Variables.

|  | Females | | | | | Males | | | | |
| --- | --- | --- | --- | --- | --- | --- | --- | --- | --- | --- |
| Variable | Sample size | R^2^ | Cohen's f | Power (1-β) | Samples per group (Power = 0.80) | Sample size | R^2^ | Cohen's f | Power (1-β) | Samples per group (Power = 0.80) |
| Bone Length (mm) | 20 | 0.320 | 0.686 | 0.710 | 8 | 21 | 0.467 | 0.936 | 0.950 | 5 |
| Tb.BV/TV (%) | 20 | 0.055 | 0.242 | 0.131 | 56 | 21 | 0.782 | 1.891 | 1.000 | 3 |
| Tb. BMD  (mgHA/cm^3^) | 20 | 0.038 | 0.198 | 0.103 | 84 | 21 | 0.804 | 2.023 | 1.000 | 3 |
| ConnD. (1/mm^3^) | 20 | 0.888 | 2.810 | 1.000 | 2 | 21 | 0.866 | 2.544 | 1.000 | 2 |
| Tb. N  (1/mm) | 20 | 0.532 | 1.067 | 0.980 | 5 | 21 | 0.888 | 2.820 | 1.000 | 2 |
| Tb. Th  (mm) | 20 | 0.176 | 0.461 | 0.375 | 17 | 21 | 0.017 | 0.130 | 0.073 | 193 |
| Tb. Sp  (mm) | 20 | 0.422 | 0.854 | 0.887 | 6 | 21 | 0.914 | 3.256 | 1.000 | 2 |
| B.Ar  (mm^2^) | 20 | 0.560 | 1.128 | 0.989 | 4 | 21 | 0.368 | 0.763 | 0.827 | 7 |
| Ma. Ar  (mm^2^) | 20 | 0.034 | 0.186 | 0.097 | 94 | 21 | 0.769 | 1.823 | 1.000 | 3 |
| Tt.Ar  (mm^2^) | 20 | 0.422 | 0.854 | 0.887 | 6 | 21 | 0.713 | 1.577 | 1.000 | 3 |
| B. Ar/Tt.Ar  (%) | 20 | 0.080 | 0.294 | 0.174 | 39 | 21 | 0.752 | 1.739 | 1.000 | 3 |
| Ct.Ar  (mm^2^) | 20 | 0.480 | 0.961 | 0.949 | 5 | 21 | 0.372 | 0.770 | 0.834 | 7 |
| Ct. Ar/Tt. Ar  (%) | 20 | 0.254 | 0.584 | 0.561 | 11 | 21 | 0.726 | 1.629 | 1.000 | 3 |
| Ct. Th  (mm) | 20 | 0.227 | 0.542 | 0.497 | 12 | 21 | 0.747 | 1.720 | 1.000 | 3 |
| Ct. TMD  (mgHA/cm^3^) | 20 | 0.771 | 1.833 | 1.000 | 3 | 21 | 0.298 | 0.651 | 0.688 | 9 |
| Ct. Po  (%) | 20 | 0.405 | 0.825 | 0.864 | 6 | 21 | 0.651 | 1.365 | 1.000 | 3 |
| C_min_  (mm) | 20 | 0.098 | 0.329 | 0.209 | 31 | 21 | 0.864 | 2.518 | 1.000 | 2 |
| pMOI  (mm^4^) | 20 | 0.495 | 0.991 | 0.960 | 5 | 21 | 0.640 | 1.332 | 0.999 | 4 |
| I_max_  (mm^4^) | 20 | 0.535 | 1.072 | 0.981 | 4 | 21 | 0.545 | 1.095 | 0.989 | 4 |
| I_min_  (mm^4^) | 20 | 0.363 | 0.755 | 0.795 | 7 | 21 | 0.729 | 1.641 | 1.000 | 3 |
| Section Modulus (mm^3^) | 20 | 0.441 | 0.888 | 0.911 | 6 | 21 | 0.566 | 1.143 | 0.994 | 4 |
| CTX1/P1NP | 26 | 0.042 | 0.209 | 0.110 | 75 | 22 | 0.130 | 0.386 | 0.302 | 23 |
| Periosteal  Calcein (%) | 20 | 0.014 | 0.121 | 0.069 | 221 | 20 | 0.150 | 0.420 | 0.317 | 20 |
| Endosteal  Calcein (%) | 20 | 0.076 | 0.287 | 0.167 | 41 | 20 | 0.164 | 0.442 | 0.348 | 185 |
| Total  Calcein (%) | 20 | 0.022 | 0.150 | 0.080 | 144 | 20 | 0.089 | 0.313 | 0.192 | 34 |
| Periosteal Alizarin (%) | 20 | 0.001 | 0.035 | 0.052 | 2675 | 20 | 0.042 | 0.208 | 0.109 | 76 |
| Endosteal Alizarin (%) | 20 | 0.122 | 0.372 | 0.257 | 25 | 20 | 0.173 | 0.457 | 0.369 | 17 |
| Total  Alizarin (%) | 20 | 0.028 | 0.171 | 0.089 | 112 | 20 | 0.001 | 0.037 | 0.052 | 2292 |
| Stiffness (N/mm) | 20 | 0.413 | 0.839 | 0.876 | 6 | 21 | 0.117 | 0.365 | 0.261 | 26 |
| Max Load  (N) | 20 | 0.131 | 0.389 | 0.277 | 23 | 21 | 0.045 | 0.217 | 0.119 | 70 |
| Fracture Load (N) | 20 | 0.161 | 0.438 | 0.342 | 18 | 21 | 0.054 | 0.238 | 0.134 | 58 |
| Energy at Fracture  (mJ) | 20 | 0.236 | 0.555 | 0.517 | 12 | 21 | 0.244 | 0.567 | 0.562 | 12 |
| Modulus (GPa) | 20 | 0.017 | 0.132 | 0.073 | 185 | 21 | 0.484 | 0.969 | 0.962 | 5 |
| Yield Stress (MPa) | 20 | 0.284 | 0.630 | 0.631 | 10 | 21 | 0.185 | 0.477 | 0.419 | 16 |
| Post Yield Strain | 20 | 0.060 | 0.254 | 0.140 | 51 | 21 | 0.033 | 0.184 | 0.098 | 97 |
| Ultimate Stress  (MPa) | 20 | 0.280 | 0.623 | 0.620 | 10 | 21 | 0.443 | 0.892 | 0.929 | 6 |
| Toughness (N/mm^2^) | 20 | 0.429 | 0.866 | 0.896 | 6 | 21 | 0.363 | 0.755 | 0.819 | 7 |
| ν_2_PO_4_^-3^: Amide III | 19 | 0.013 | 0.115 | 0.066 | 243 | 21 | 0.324 | 0.692 | 0.743 | 8 |
| CO_3_^-2^:  ν_1_PO_4_^-3^ | 19 | 0.009 | 0.096 | 0.061 | 351 | 21 | 0.016 | 0.129 | 0.073 | 195 |
| Crystallinity (cm^-1^) | 19 | 0.036 | 0.194 | 0.098 | 87 | 21 | 0.015 | 0.123 | 0.071 | 215 |
| I1670:I1640 | 19 | 0.074 | 0.283 | 0.157 | 42 | 21 | 0.388 | 0.797 | 0.860 | 7 |
| I1670:I1610 | 19 | 0.086 | 0.306 | 0.177 | 36 | 21 | 0.404 | 0.824 | 0.883 | 6 |
| I1670:I1690 | 19 | 0.121 | 0.372 | 0.243 | 25 | 21 | 0.005 | 0.067 | 0.056 | 712 |
| K_c,init_  (MPa m^1/2^) | 16 | 0.273 | 0.613 | 0.483 | 10 | 14 | 0.250 | 0.577 | 0.373 | 11 |
| K_c,max_  (MPa m^1/2^) | 19 | 0.414 | 0.840 | 0.854 | 6 | 19 | 0.159 | 0.435 | 0.319 | 19 |

S1. Load-displacement curve cropped near first failure demonstrating strain hardening.

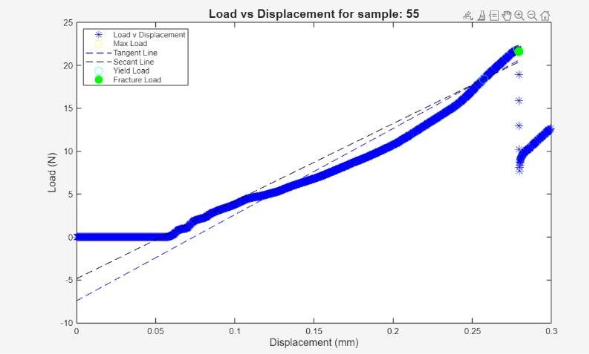

S2. **A**. Representative quantitative histomorphology images of cortical diaphysis for groups showing calcein (green, -10d) and alizarin (red, -3d) labeling. **B**. The CTX1/P1NP ratio does not significantly change between groups. **C**. The total percentage of cortical surface labeled 10 days prior to euthanasia does not change with the experimental groups; however, 7-mo males exhibited high variance that may affect interpretation. **D**. The total labeled cortical surface is not distinct between the observed groups.

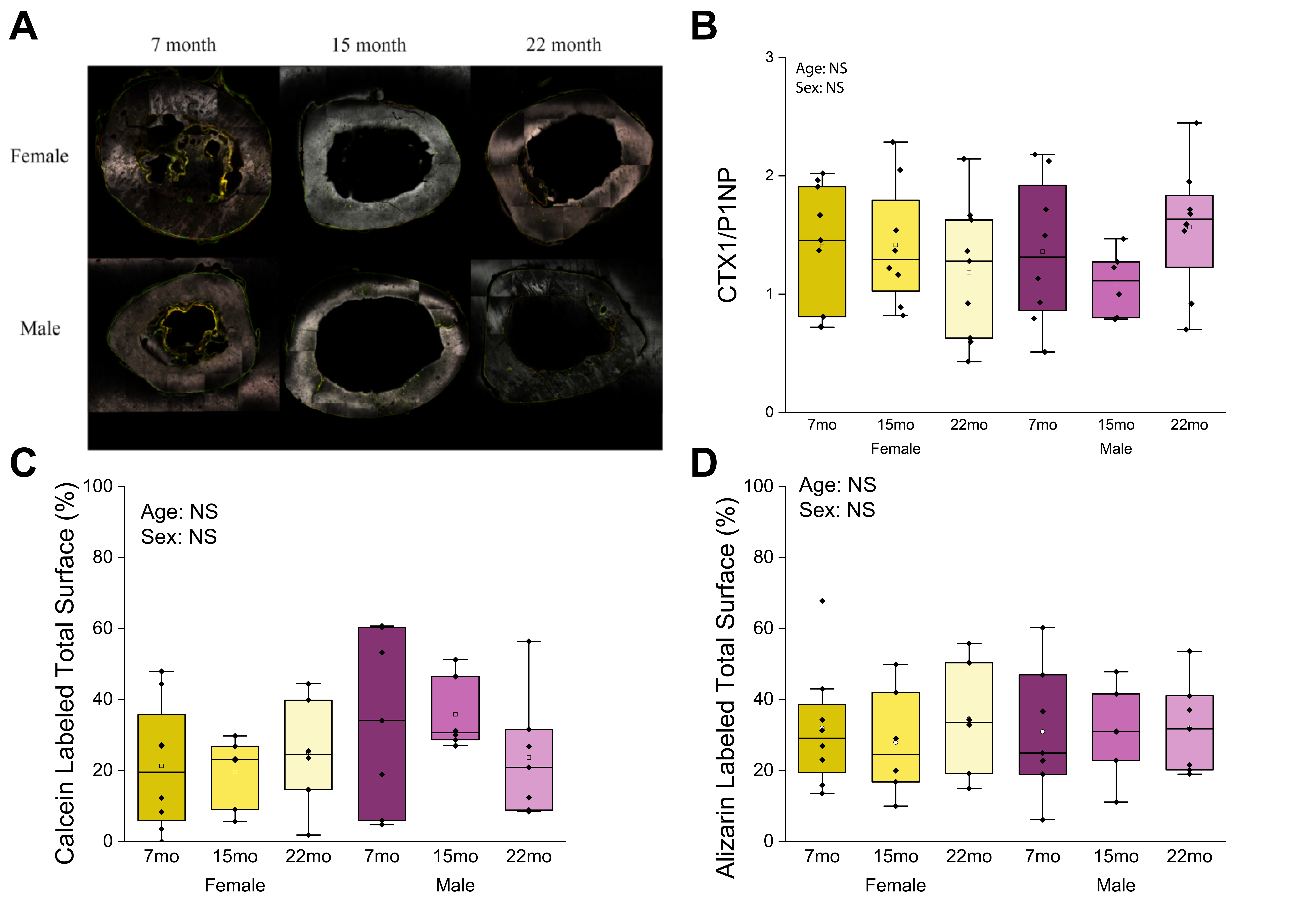

S3. Stress-strain curves with group membership.

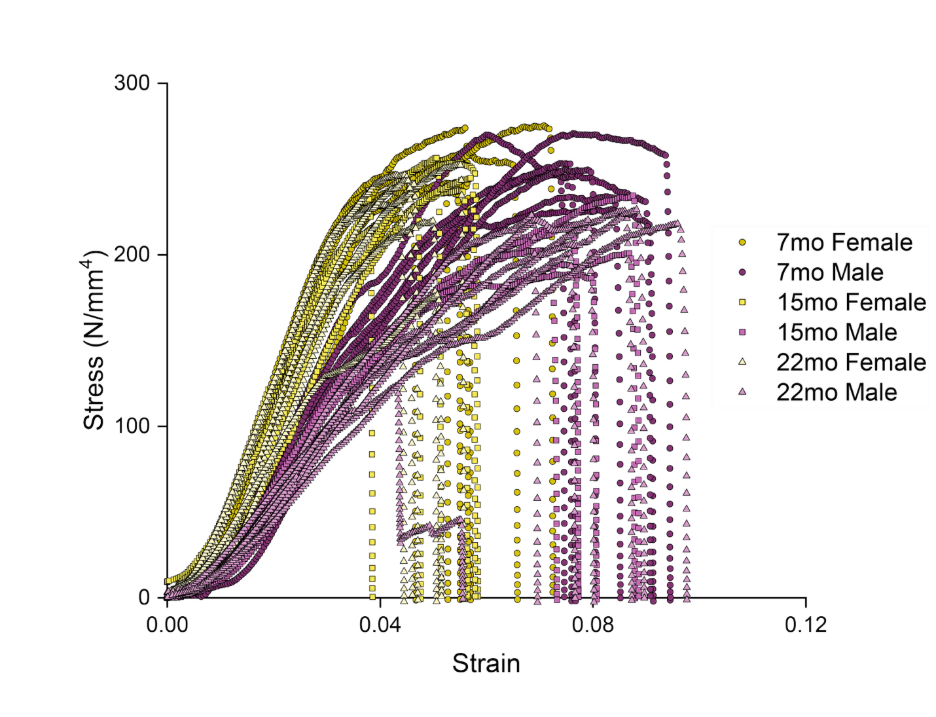

S4. Load-displacement curves with group membership.

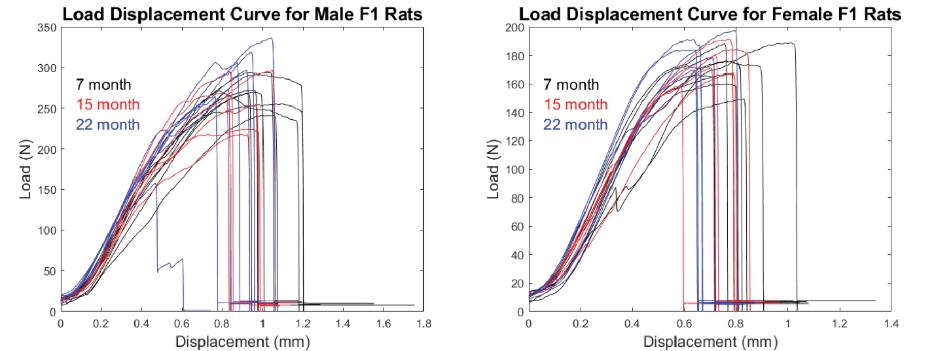

S5. Box plot of K_c,init_ for groups.

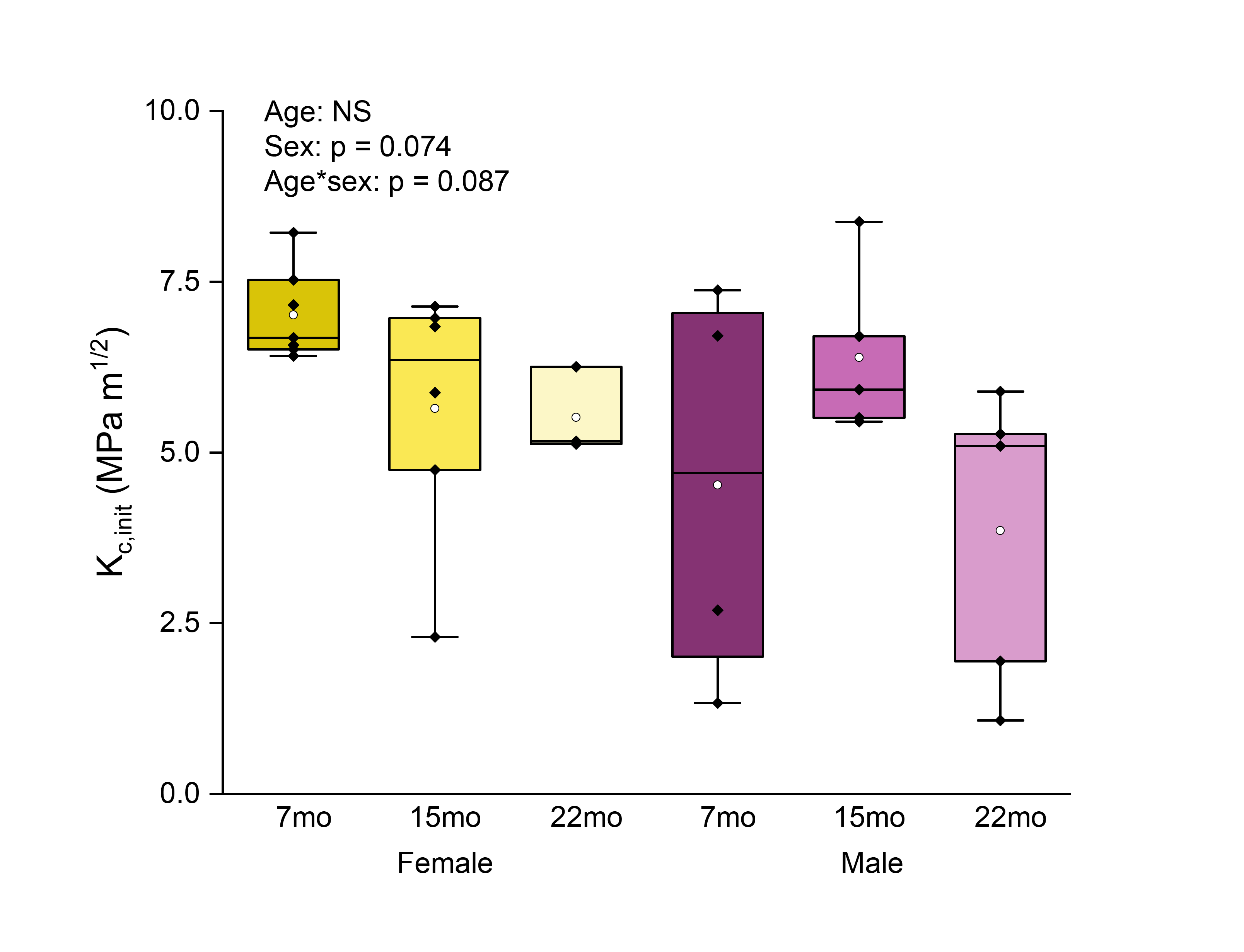

S6. Representative image from fracture toughness testing.

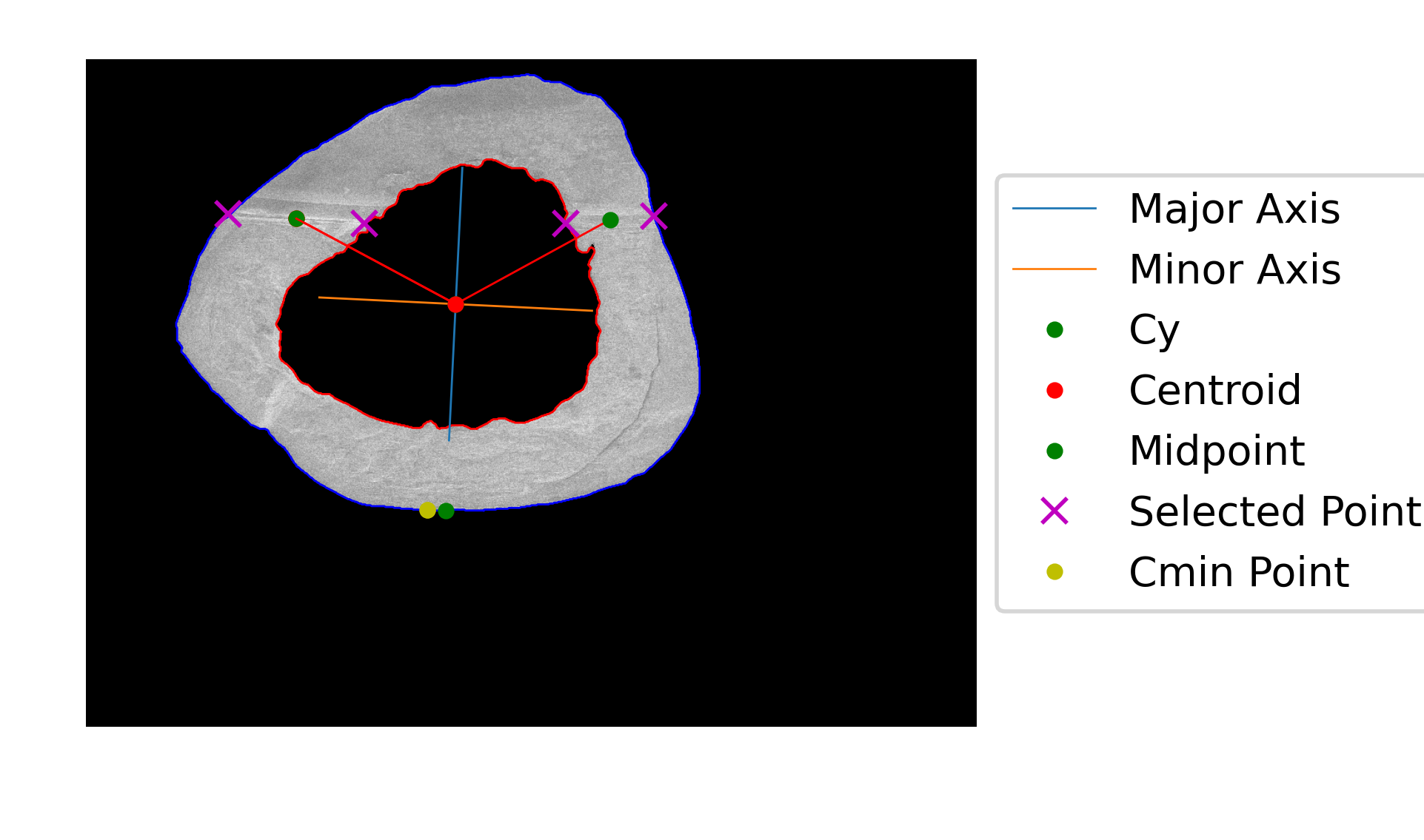

S7: PCA plot of bone quality measure with identity mapped.

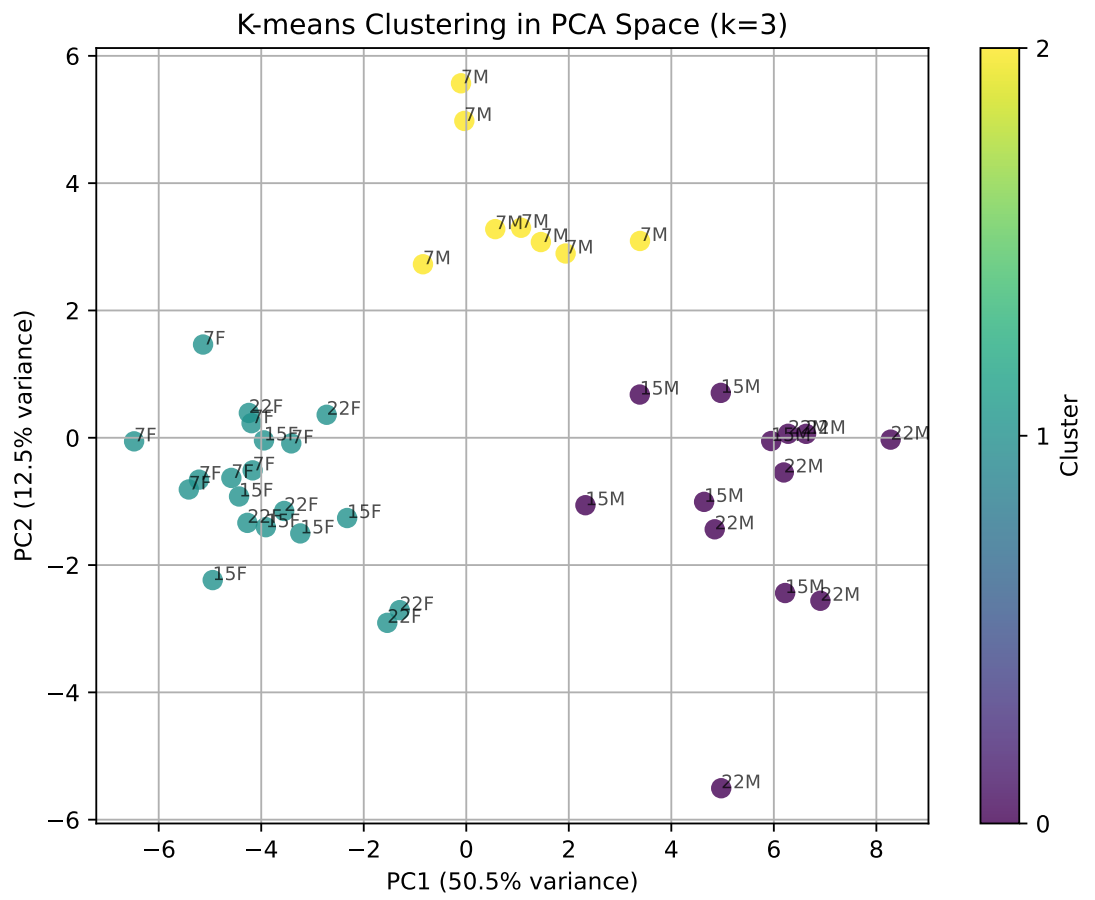

ST2. PCA loading table for PC1, PC2, and PC3.

| Variable | PC1 | PC2 | PC3 |
| --- | --- | --- | --- |
| Tt. Ar (mm^2^) | 0.225985 | 0.015006 | -0.00049 |
| I_min_(mm^4^) | 0.225625 | 0.008878 | 0.002765 |
| pMOI(mm^4^) | 0.224491 | 0.03543 | 0.015682 |
| Section Modulus (mm^3^) | 0.224386 | 0.048679 | 0.001148 |
| I_max_(mm^4^) | 0.222049 | 0.054362 | 0.024743 |
| Ma. Ar (mm^2^) | 0.220217 | -0.07865 | -0.00865 |
| C_min_ (mm) | 0.219939 | -0.03878 | -0.02986 |
| Bone Length (mm) | 0.215556 | 0.044912 | -0.01639 |
| Ct. Ar (mm^2^) | 0.213963 | 0.129636 | -0.04835 |
| B. Ar (mm^2^) | 0.208313 | 0.128921 | 0.009267 |
| Modulus (GPa) | -0.20094 | -0.04087 | 0.069671 |
| Tb. N (1/mm) | -0.19905 | 0.169444 | 0.0793 |
| Tb. Sp (mm) | 0.198305 | -0.18756 | -0.06956 |
| Fracture Load (N) | 0.19421 | 0.192145 | 0.068479 |
| Max load (N) | 0.190391 | 0.220155 | 0.049703 |
| B. Ar/Tt. Ar (%) | -0.18109 | 0.177209 | 0.059216 |
| Tb. BMD (mgHA/cm^3^) | -0.17639 | 0.146391 | 0.164851 |
| Tb. BV/TV (%) | -0.1762 | 0.153239 | 0.169304 |
| Stiffness (N/mm) | 0.176156 | 0.052868 | 0.003631 |
| ConnD. (1/mm^3^) | -0.1686 | 0.194735 | 0.001768 |
| Ct. Ar/Tt. Ar (%) | -0.16257 | 0.227096 | -0.09613 |
| Yield stress (MPa) | -0.1511 | 0.084622 | 0.175482 |
| Energy at fracture (mJ) | 0.143493 | 0.332523 | 0.015327 |
| Post yield strain | 0.139019 | 0.213883 | -0.07497 |
| Ultimate Stress (MPa) | -0.12818 | 0.279141 | 0.091607 |
| I1670: I1610 | 0.107004 | -0.09995 | 0.300108 |
| Carbonate: phosphate | -0.10246 | -0.03349 | -0.41215 |
| Crystallinity | 0.08685 | 0.026861 | 0.394708 |
| Ct. Po (%) | 0.084174 | -0.17763 | 0.055598 |
| Toughness (N/mm^2^) | 0.080023 | 0.379799 | 0.000417 |
| I1670: I1640 | 0.068182 | -0.10153 | 0.381556 |
| v2phos: Amide III | -0.06816 | -0.04544 | -0.20633 |
| Kc,init (MPa m^1/2^) | -0.05927 | 0.004052 | 0.201989 |
| Kc,max (MPa m^1/2^) | -0.0576 | 0.0024 | 0.234611 |
| Ct. Th (mm) | 0.042226 | 0.357432 | -0.14141 |
| Tb. Th (mm) | 0.034616 | 0.103048 | 0.233709 |
| I1670: I1690 | -0.00684 | 0.113965 | -0.27681 |
| Ct. TMD (mgHA/cm^3^) | -0.00514 | -0.194 | 0.068066 |

S8. Cortical metabolome (A) PCA, (B) PLS-DA, (C) heatmap with all features, and volcano plots for significantly different metabolomes [(D) 7-mo males vs 22-mo males, (E) 7-mo females vs 22-mo females, (F) 22-mo males vs 22-mo females].

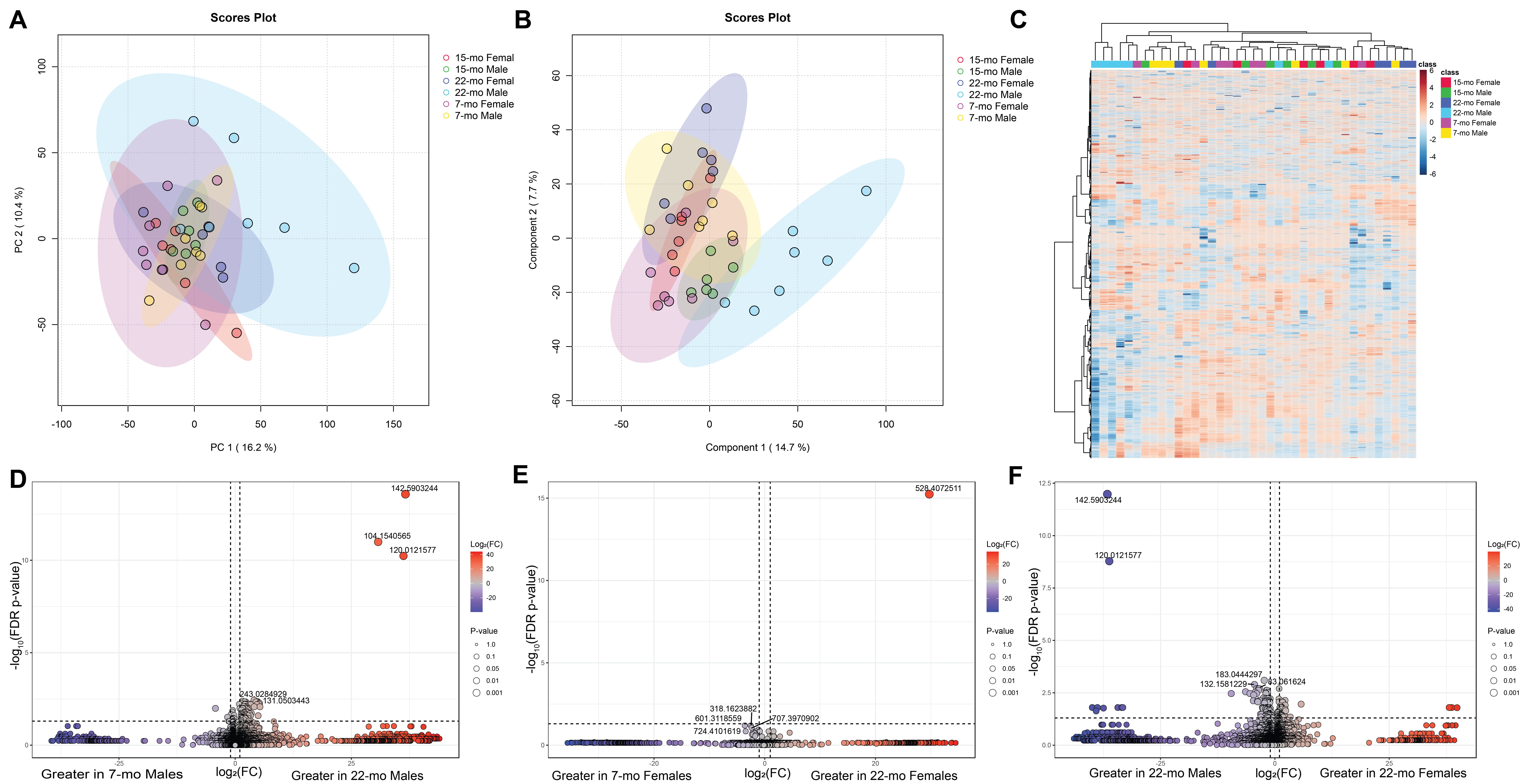

ST3. Pseudovariable composition table for skeletal aging in long bones.

| Pseudovariable | Compositional Variables |
| --- | --- |
| Bone Quantity | Tb. BMD, Tb. TMD, Tb. BV/TV, Ct. BMD, Ct. TMD, Ct. BV/TV, Ct. Ar/Tt. Ar |
| Bone Quality | Conn. D, Tb. Th., Tb. N, Ct. Po, Ct. Th, ν1PO4: Amide I, ν_1_PO_4_: Amide III, ν_2_PO_4_: Amide III, ν _1_PO_4_: proline, crystallinity, carbonate: phosphate, I1670: I1610, I1670: I1640, I1670: I1690, bound water, pyridinoline per collagen, deoxypyridinoline per collagen, pentosidine per collagen |
| Fracture Resistance | Ultimate stress, maximum load, energy-to-fracture, toughness, K_c,init_, K_c,max_, yield moment, yield stress, elastic modulus, post yield deformation, post yield strain, strength-strain index |
| Health in Aging | ln(median lifespan), spontaneous tumorigenesis |

Supplemental references.

1. Bloomfield SA, Hogan HA, Delp MD. Decreases in Bone Blood Flow and Bone Material Properties in Aging Fischer-344 Rats. Clinical Orthopaedics and Related Research (1976-2007). 2002;396
2. Burstein AH, Reilly DT, Martens M. Aging of bone tissue: mechanical properties. JBJS. 1976;58(1)
3. Byberg L, Gedeborg R, Cars T, et al. Prediction of fracture risk in men: A cohort study*. Journal of Bone and Mineral Research. 2012;27(4):797-807. doi:10.1002/jbmr.1498
4. Cirovic A, Jadzic J, Djukic D, et al. Increased Cortical Porosity, Reduced Cortical Thickness, and Reduced Trabecular and Cortical Microhardness of the Superolateral Femoral Neck Confer the Increased Hip Fracture Risk in Individuals with Type 2 Diabetes. Calcified Tissue International. 2022/11/01 2022;111(5):457-465. doi:10.1007/s00223-022-01007-6
5. ENVIGO. BALB/c Bagg's Albino. <https://insights.envigo.com/hubfs/resources/data-sheets/envigo-49-balbc-letter_screen.pdf>
6. Ferguson VL, Ayers RA, Bateman TA, Simske SJ. Bone development and age-related bone loss in male C57BL/6J mice. Bone. 2003/09/01/ 2003;33(3):387-398. doi:<https://doi.org/10.1016/S8756-3282(03)00199-6>
7. Francisco JI, Yu Y, Oliver RA, Walsh WR. Relationship between age, skeletal site, and time post-ovariectomy on bone mineral and trabecular microarchitecture in rats. Journal of Orthopaedic Research. 2011/02/01 2011;29(2):189-196. doi:<https://doi.org/10.1002/jor.21217>
8. Fukuda S, Iida H. Age-Related Changes in Bone Mineral Density, Cross-Sectional Area and the Strength of Long Bones in the Hind Limbs and First Lumbar Vertebra in Female Wistar Rats. Journal of Veterinary Medical Science. 2004;66(7):755-760. doi:10.1292/jvms.66.755
9. Gerstenfeld LC, McLean J, Healey DS, et al. Genetic variation in the structural pattern of osteoclast activity during post-natal growth of mouse femora. Bone. 2010/06/01/ 2010;46(6):1546-1554. doi:<https://doi.org/10.1016/j.bone.2010.02.016>
10. Glatt V, Canalis E, Stadmeyer L, Bouxsein ML. Age-Related Changes in Trabecular Architecture Differ in Female and Male C57BL/6J Mice. Journal of Bone and Mineral Research. 2007/08/01 2007;22(8):1197-1207. doi:<https://doi.org/10.1359/jbmr.070507>
11. Halloran BP, Ferguson VL, Simske SJ, Burghardt A, Venton LL, Majumdar S. Changes in Bone Structure and Mass With Advancing Age in the Male C57BL/6J Mouse. Journal of Bone and Mineral Research. 2002/06/01 2002;17(6):1044-1050. doi:<https://doi.org/10.1359/jbmr.2002.17.6.1044>
12. Hernandez CJ, Stein EM, Donnelly E. Impaired Bone Matrix: The Key to Fragility in Type 2 Diabetes? The Journal of Clinical Endocrinology & Metabolism. 2021;106(7):e2825-e2827. doi:10.1210/clinem/dgab150
13. Holmes DJ. F344BNF1 and BNF344F1 Hybrid Rats. Science of Aging Knowledge Environment. 2004/11/17 2004;2004(46):as4-as4. doi:10.1126/sageke.2004.46.as4
14. Iida H, Fukuda S. Age-Related Changes in Bone Mineral Density, Cross-Sectional Area and Strength at Different Skeletal Sites in Male Rats. Journal of Veterinary Medical Science. 2002;64(1):29-34. doi:10.1292/jvms.64.29
15. Jast J, Jasiuk I. Age-related changes in the 3D hierarchical structure of rat tibia cortical bone characterized by high-resolution micro-CT. Journal of Applied Physiology. 2013/04/01 2013;114(7):923-933. doi:10.1152/japplphysiol.00948.2011
16. Macdonald HM, Nishiyama KK, Kang J, Hanley DA, Boyd SK. Age-related patterns of trabecular and cortical bone loss differ between sexes and skeletal sites: A population-based HR-pQCT study. Journal of Bone and Mineral Research. 2011/01/01 2011;26(1):50-62. doi:<https://doi.org/10.1002/jbmr.171>
17. McCalden RW, McGeough JA, Barker MB, Court-Brown CM. Age-related changes in the tensile properties of cortical bone. The relative importance of changes in porosity, mineralization, and microstructure. JBJS. 1993;75(8)
18. Mumtaz H, Dallas M, Begonia M, et al. Age-related and sex-specific effects on architectural properties and biomechanical response of the C57BL/6N mouse femur, tibia and ulna. Bone Reports. 2020/06/01/ 2020;12:100266. doi:<https://doi.org/10.1016/j.bonr.2020.100266>
19. Nnakwe NE. The effect of aging on bone composition of female Fischer 344 rats. Mechanisms of Ageing and Development. 1995/11/24/ 1995;85(2):125-131. doi:<https://doi.org/10.1016/0047-6374(95)01668-6>
20. Nyman JS, Roy A, Acuna RL, et al. Age-related effect on the concentration of collagen crosslinks in human osteonal and interstitial bone tissue. Bone. 2006/12/01/ 2006;39(6):1210-1217. doi:<https://doi.org/10.1016/j.bone.2006.06.026>
21. Nyman JS, Roy A, Tyler JH, Acuna RL, Gayle HJ, Wang X. Age-related factors affecting the postyield energy dissipation of human cortical bone. Journal of Orthopaedic Research. 2007/05/01 2007;25(5):646-655. doi:<https://doi.org/10.1002/jor.20337>
22. Osterhoff G, Morgan EF, Shefelbine SJ, Karim L, McNamara LM, Augat P. Bone mechanical properties and changes with osteoporosis. Injury. 2016/06/01/ 2016;47:S11-S20. doi:<https://doi.org/10.1016/S0020-1383(16)47003-8>
23. Papageorgiou M, Föger-Samwald U, Wahl K, Kerschan-Schindl K, Pietschmann P. Age- and Strain-Related Differences in Bone Microstructure and Body Composition During Development in Inbred Male Mouse Strains. Calcified Tissue International. 2020/04/01 2020;106(4):431-443. doi:10.1007/s00223-019-00652-8
24. Perrien DS, Akel NS, Dupont-Versteegden EE, et al. Aging alters the skeletal response to disuse in the rat. American Journal of Physiology-Regulatory, Integrative and Comparative Physiology. 2007/02/01 2007;292(2):R988-R996. doi:10.1152/ajpregu.00302.2006
25. Pietschmann P, Skalicky M, Kneissel M, et al. Bone structure and metabolism in a rodent model of male senile osteoporosis. Experimental Gerontology. 2007/11/01/ 2007;42(11):1099-1108. doi:<https://doi.org/10.1016/j.exger.2007.08.008>
26. Price C, Herman BC, Lufkin T, Goldman HM, Jepsen KJ. Genetic Variation in Bone Growth Patterns Defines Adult Mouse Bone Fragility. Journal of Bone and Mineral Research. 2005/11/01 2005;20(11):1983-1991. doi:<https://doi.org/10.1359/JBMR.050707>
27. Puelker SM, Ribeiro de Castro SR, de Souza RR, Maifrino LBM, Nucci RAB, Sitta MdC. Age-Related Effects on Right Femoral Bone of Male Wistar Rats: A Morphometric and Biomechanical Study. Journal of Health and Allied Sciences NU. 2021/06/01 2021;12(01):67-70. doi:10.1055/s-0041-1730107
28. Riggs BL, Melton Iii LJ, Robb RA, et al. Population-Based Study of Age and Sex Differences in Bone Volumetric Density, Size, Geometry, and Structure at Different Skeletal Sites. Journal of Bone and Mineral Research. 2004;19(12):1945-1954. doi:<https://doi.org/10.1359/jbmr.040916>
29. Sampson HW. Effect of Alcohol Consumption on Adult and Aged Bone: A Histomorphometric Study of the Rat Animal Model. Alcoholism: Clinical and Experimental Research. 1998/12/01 1998;22(9):2029-2034. doi:<https://doi.org/10.1111/j.1530-0277.1998.tb05912.x>
30. Silbermann M, Safadi M, Schapira D, Leichter I, Steinberg R. Structural and compositional changes in aging bone: osteopenia in lumbar vertebrae of Wistar female rats. Scanning Microscopy. 1989;(0891-7035 (Print))
31. Silbermann M, Weiss A, Reznick AZ, Eilam Y, Szydel N, Gershon D. Age-related trend for osteopenia in femurs of female C57BL/6 mice. Compr Gerontol A. 1987/02// 1987;1(1):45-51.
32. Singulani MP, Stringhetta-Garcia CT, Santos LF, et al. Effects of strength training on osteogenic differentiation and bone strength in aging female Wistar rats. Scientific Reports. 2017/02/17 2017;7(1):42878. doi:10.1038/srep42878
33. Sprott RL. Development of animal models of aging at the national institute on aging. Neurobiology of Aging. 1991/11/01/ 1991;12(6):635-638. doi:<https://doi.org/10.1016/0197-4580(91)90113-X>
34. Zhang H, Bethel CS, Smittkamp SE, Stanford JA. Age-related changes in orolingual motor function in F344 vs F344/BN rats. Physiology & Behavior. 2008/02/27/ 2008;93(3):461-466. doi:<https://doi.org/10.1016/j.physbeh.2007.10.004>
35. Zhang R, Gong H, Zhu D, Ma R, Fang J, Fan Y. Multi-level femoral morphology and mechanical properties of rats of different ages. Bone. 2015/07/01/ 2015;76:76-87. doi:<https://doi.org/10.1016/j.bone.2015.03.022>
36. Zioupos P, Currey JD, Hamer AJ. The role of collagen in the declining mechanical properties of aging human cortical bone. Journal of Biomedical Materials Research. 1999/05/01 1999;45(2):108-116. doi:https://doi.org/10.1002/(SICI)1097-4636(199905)45:2%3C108::AID-JBM5%3E3.0.CO;2-A
37. Vogel HG. Influence of maturation and aging on mechanical and biochemical properties of connective tissue in rats. Mechanisms of Ageing and Development. 1980/11/01/ 1980;14(3):283-292. doi:<https://doi.org/10.1016/0047-6374(80)90002-0>
